# A predictive processing model of episodic memory and time perception

**DOI:** 10.1101/2020.02.17.953133

**Authors:** Zafeirios Fountas, Anastasia Sylaidi, Kyriacos Nikiforou, Anil K. Seth, Murray Shanahan, Warrick Roseboom

**Affiliations:** Wellcome Centre for Human Neuroimaging, Institute of Neurology, University College London, London, UK; Emotech Labs, London, UK; Spike AI Research Labs, London, UK; Department of Computing, Imperial College London, London, UK; Department of Informatics and Sackler Centre for Consciousness Science, University of Sussex, Sussex, UK; Canadian Institute for Advanced Research (CIFAR) Program on Brain, Mind, and Consciousness, Toronto, Ontario, Canada

## Abstract

Human perception and experience of time is strongly influenced by ongoing stimulation, memory of past experiences, and required task context. When paying attention to time, time experience seems to expand; when distracted, it seems to contract. When considering time based on memory, the experience may be different than in the moment, exemplified by sayings like “time flies when you’re having fun”. Experience of time also depends on the content of perceptual experience – rapidly changing or complex perceptual scenes seem longer in duration than less dynamic ones. The complexity of interactions between attention, memory, and perceptual stimulation is a likely reason that an overarching theory of time perception has been difficult to achieve. Here, we introduce a model of perceptual processing and episodic memory that makes use of hierarchical predictive coding, short-term plasticity, spatio-temporal attention, and episodic memory formation and recall, and apply this model to the problem of human time perception. In an experiment with ~ 13, 000 human participants we investigated the effects of memory, cognitive load, and stimulus content on duration reports of dynamic natural scenes up to ~ 1 minute long. Using our model to generate duration estimates, we compared human and model performance. Model-based estimates replicated key qualitative biases, including differences by cognitive load (attention), scene type (stimulation), and whether the judgement was made based on current or remembered experience (memory). Our work provides a comprehensive model of human time perception and a foundation for exploring the computational basis of episodic memory within a hierarchical predictive coding framework.

**Author summary:** Experience of the duration of present or past events is a central aspect of human experience, the underlying mechanisms of which are not yet fully understood. In this work, we combine insights from machine learning and neuroscience to propose a combination of mathematical models that replicate human perceptual processing, long-term memory, attention, and duration perception. Our computational implementation of this framework can process information from video clips of ordinary life scenes, record and recall important events, and report the duration of these clips. To assess the validity of our proposal, we conducted an experiment with ~ 13, 000 human participants. Each was shown a video between 1-64 seconds long and reported how long they believed it was. Reports of duration by our computational model qualitatively matched these human reports, made about the exact same videos. This was true regardless of the video content, whether time was actively judged or based on memory of the video, or whether the participants focused on a single task or were distracted - all factors known to influence human time perception. Our work provides the first model of human duration perception to incorporate these diverse and complex factors and provides a basis to probe the deep links between memory and time in human experience.

## Introduction

The ability to estimate temporal properties of the world, such as how much time has elapsed since you started reading this paper, is key to complex cognition, planning, and behaviour. Human perception of time is affected by many factors; experience on the scale of seconds is prominently influenced by the content, complexity, and rate of change of experience [1–4]. The influence of stimulus complexity and rate of change can be seen in cases driven by very basic stimulus properties, such as the temporal frequency [5, 6], and can also be seen in very coarse, abstract differences in content. For example, human participants report videos of scenes filmed in a busy city as longer than those filmed walking around in the countryside, with both reported as longer than scenes of a quiet office [4, 7]. Time perception is also influenced by whether attention can and is being directed toward the task of monitoring time [8–10]. Specifically, increasing cognitive load (by requiring concurrent attention to additional tasks beyond tracking time) has been reported to decrease the apparent duration of an interval when a person is aware that attention to time is required for a given task (prospective time), while apparent duration is increased by cognitive load when reflecting on the duration of an interval after it has occurred (retrospective time; comprehensively reviewed in [10] and Figure 1). Differences in time perception based on this interaction between cognitive load and prospective versus retrospective duration judgements have led to suggestions that different mechanisms underlie the different scenarios: when actively attending to time (prospective time) the process is largely “attention” driven; when reflecting on a period of time after it has occurred (retrospective time) the process is largely driven by “memory” [8, 10–12]. Reflecting these different approaches, models of time perception and memory have developed largely independently since diverging several decades ago (approach in [13] versus [14], for example), precluding a unified account of time perception.

**Fig 1.**
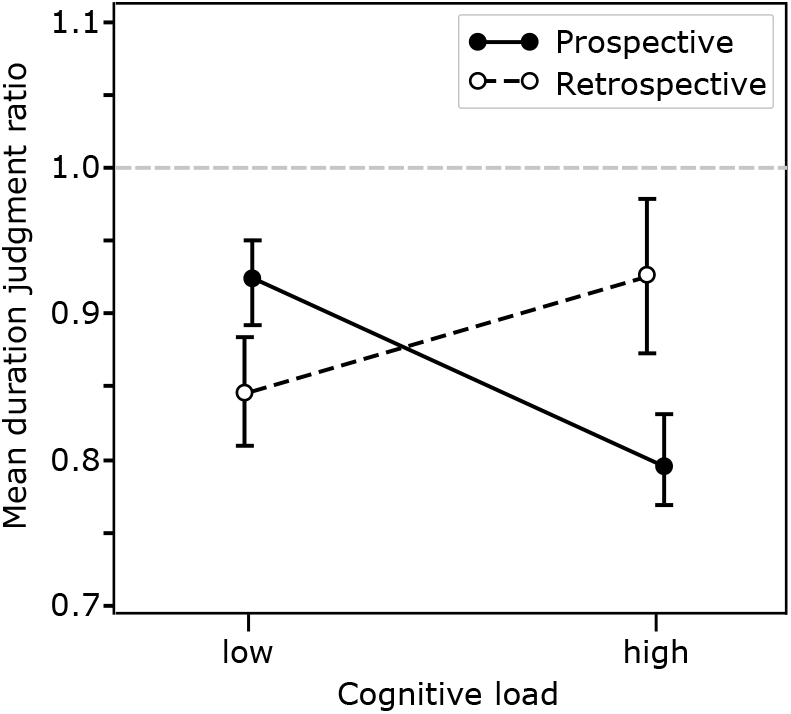
Relationship between prospective and retrospective duration judgments and their interaction with cognitive load (adapted from [10]). Duration judgement ratio is reported duration divided by physical duration. Error bars indicate standard error of the mean. Increasing cognitive load decreases reported duration in a prospective timing task, but increases reported duration in a retrospective timing task.

Here we present a model of human perceptual processing, memory, and time perception. Through this model, the influences of stimulus, attention, and content of memory on human time estimation can be integrated, via the two intertwined processes of memory formation and recall, providing a single overarching understanding of their relationship. We validate model performance in the domain of time perception by showing that model-based estimates of the duration of naturalistic videos between 1 and 64 seconds match duration reports obtained from ~ 13, 000 human participants (*n* = 6, 977 for the main results after removing all outliers) regarding the same videos for each of the key manipulations: knowingly tracking time (prospective judgements) or judging it from memory (retrospective judgements), attending to only a single or multiple concurrent tasks (cognitive load), and for different types of natural stimulation (busy city scenes, less busy scenes on a university campus or leafy surrounds, or in a quiet office of cafe).

### Time perception via perceptual classification and long-term memory

The foundation of the present model rests upon the proposal that the primary function of human perception is the classification of objects and events in the world and that time can be estimated simply by tracking the behaviour of neural networks involved in perceptual classification. Using deep convolutional neural networks (CNN) as a model of biological perceptual classification has become common in recent years (see [15] for review). CNNs have a hierarchical structure both inspired by and broadly resembling the human/primate visual cortex [16–18], and have been shown to perform well in visual classification tasks [19, 20]. While reasonable discussion continues about the extent of similarity between biological classification networks and specific artificial network architectures [21, 22], the constellation of features exhibited by CNNs has been shown to have value for investigating the bases of human perception and cognition [15, 23].

In support of the proposal that time can be estimated on the basis of activity in perceptual classification networks, a recent study [4] demonstrated that by simply tracking salient events in the activity of a CNN while it processed videos of natural scenes (from 1-64 seconds in duration) it was possible to generate a basis for prospective estimates of duration. In that study, model-based duration estimates replicated key features of human estimates for the exact same videos, including biases related to scene type (busy city scenes estimated as longer than quiet office scenes, for example; see also [7, 24]). This model displayed conceptual similarities with the predictive processing account of perception wherein perception is proposed to operate as a function of both sensory predictions and current sensory stimulation, with perceptual content understood as the brain’s “best guess” (Bayesian posterior) of the causes of current sensory input given the prior expectations or predictions [25–29].

However, this previous model [4] had only very simple “memory” that marked the occurrence of salient events in perception. Because this model lacks the ability to form content-specific memories of an episode, it is impossible to use it to estimate time in *retrospect*, where the relevant content of memory (events that have occurred - taking a bus; seeing a cow) must be used to form an estimate. In order to resolve how human retrospective judgements of time might be accomplished, we extended on the core aspects of this previous model to provide the ability to record episodes of perceptual content as memory.

To understand the relation between memory and time in perception we first need to define a theoretical model of long-term memory. Explicit long-term memory is often divided into “semantic” and “episodic”, reflecting the difference between “knowing” and “remembering” [30]. Whereas semantic memory maintains generalized (statistical) information about different concepts in the world, including taxonomic (i.e. similarity-based) and thematic (i.e. contiguity-based) relations [31], episodic memory stores individual events, including specific temporal contiguity relations between them [32]. These two memory systems are believed to be based on highly interdependent mechanisms, during both formation and recall [33]. In what follows, we first present a novel Bayesian model of episodic and semantic memory, defined under the predictive processing framework. We then focus on the proposed mechanisms for basic memory operations and their role in generating the inner sense of duration. Next, we compare the performance of this proposed model against human reports of duration in the order of seconds (up to 64 seconds). Finally, we discuss the relationship between our new model and prominent work on the neural foundations of episodic memory in event segmentation and temporal hierarchy in the brain.

## Theoretical framework

### Predictive processing neural architecture for perception

According to (Bayesian) predictive processing theories, the brain is constantly updating its internal representations of the state of the world with new sensory information 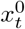 (at time *t*) through the process of hierarchical probabilistic inference [25, 28, 34, 35]. The resulting ‘top-down’ generative model is used to predict new states and simulate the outcome of prospective actions [36]. Comparing predicted against actual sensory signals gives rise to prediction errors 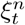 which, in most instantiations of predictive processing, are assumed to flow in a ‘bottom-up’ direction from the sensory periphery (*n* = 0), towards higher hierarchical levels *n* ∈ {1, 2,..}. Numerous connections with neurophysiology have been made over the years, including proposals viewing predictive processing as the fundamental function of canonical microcircuits in the cortex [37], or as the result of the interplay between competing oscillatory bands [38, 39], as well as providing theoretical explanations for a wide range of cognitive and perceptual effects such as hallucinations [40], autism [41] and schizophrenia [42]. In addition, this framework is uniquely qualified for studying the relation between perceptual change and human perception of time [4], as it provides a working explanation of the complex interplay between prior beliefs, sensory-based information and learning.

In this study, we define a predictive processing model that relies on feed-forward deep neural networks for the bottom-up flow of information (see *inference model* in Figure 2) and a novel stochastic process to account for the top-down flow of predictions (see *generative model* in the same figure). Unlike famous approaches that use neural networks to implement inference via amortization [43, 44], here neural activations of each hierarchical layer *n* represent single samples of the random variable 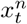, rather than its distribution parameters. Hence, existing, pre-trained, feed-forward neural networks can be used to propagate bottom-up sensory signals or, in this case, to approximate the inference 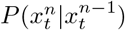.

**Fig 2.**
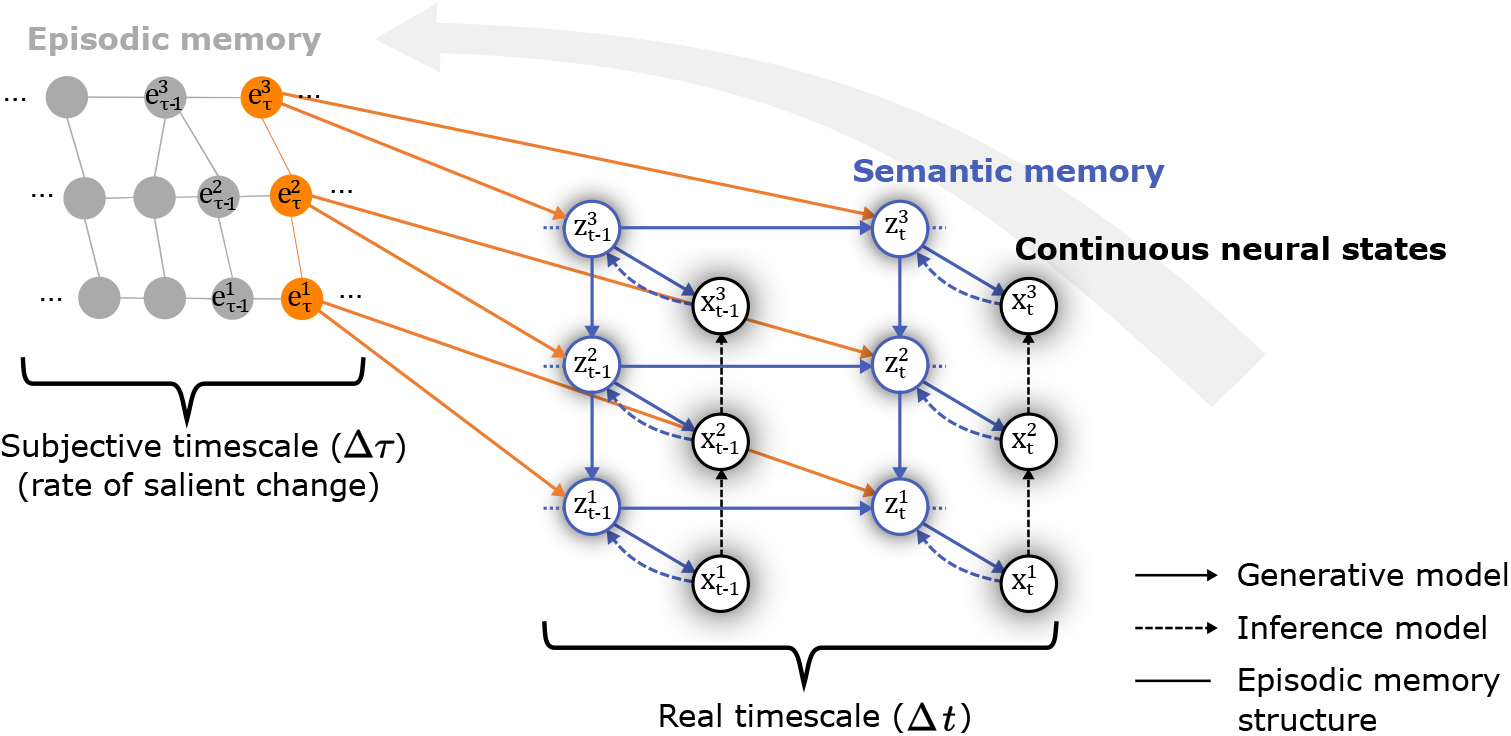
Simplified probabilistic graphical representation of the proposed predictive processing model described in the text, highlighting the distinction between episodic and semantic memory systems. The random variables 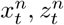 correspond to different hierarchical representations of the input, continuous and categorical respectively. All solid-line arrows represent conditional dependencies while, in particular, orange arrows highlight dependencies between variables with distributions that evolve over different time scales. For a full graphical representation of the model equations see S1 Fig.

The structure of the generative model (solid arrows in Figure 2 and S1 Fig) can be best described using the three major perceptual principles that it was designed to account for. First, humans are able to segregate raw sensory information into different categories of concepts based on similarity (often called taxonomic relations) and access these categories linguistically, answering for instance questions such as *what does an object X look like?* Likewise, the model presented here generates categorical hierarchical latent representations of the sensory signals it receives, in the form of probability distributions, that classify the current (continuous) neural states and create new predictions. Second, humans can associate sensory experiences with context and with other experiences over time (often called contiguity relations), and answer questions such as *do I expect to see object X under the current circumstances?* Our model also maintains state transition statistics, taking into account the hierarchy of the categorical representations and recent experiences. Finally, humans are able to maintain the general concept of an object X, without being continuously surprised after seeing a particular instance (e.g. a paper sketch) of this object. To account for this ability, each categorical representation in the model consists of a slowly-changing baseline distribution as well as a second, short-term bias.

To implement the first principle, the (continuous) random variable 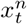 follows a mixture of Gaussians, where the parameters of the individual Gaussian components 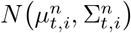 update over time with recursive Bayesian estimation^1^ At each simulation time-step and layer *n*, a single component is selected, then used to sample a prediction 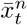, and finally updated according to the prediction error 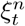. Hence,

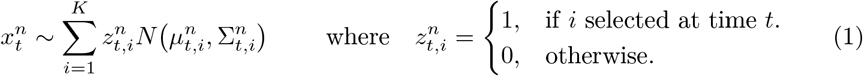

The selected component is indicated by the (discrete) random variable 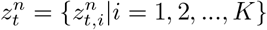 which allows the categorization of the model’s (both sensory and inner) states. Given a current observation 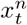, the value of 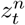 can be inferred from arg max_*i*_ 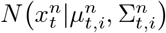. In addition, to cover the second principle, the tuple 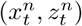 is treated as a partially-observable hierarchical hidden Markov model over time *t*, with a discrete state-space and continuous observations. Apart from the previous and current higher order states, which are used to satisfy the Markovian property, the categorical variable 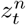 is also conditioned on the *last surprising state* that the system recorded. For now, we will assign the symbol 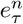 to represent this state and later in the text we will see that 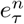 corresponds to the last recorded event in episodic memory. Taking all together, 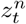 can be drawn from 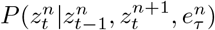. Finally, to address the third principle we propose an extension to the classical Kalman filter [45] that takes into account the differences between short- and long-term learning. Based on this proposal, each Gaussian component maintains two states, a short-term state 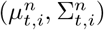 and a baseline state 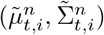. Hence, the update process of each component becomes two-fold. The short-term states update according to

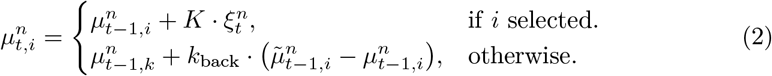

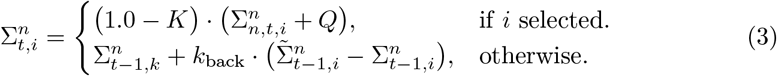

where 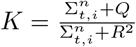 is the Kalman gain, *Q* and *R* are free noise parameters and *k*_back_ is a parameter determining how quickly the short-term state converges back to its baseline, in the absence of new information. The update of the baseline states is given by

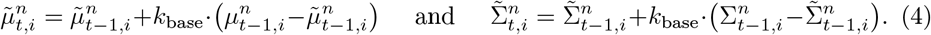

where *k*_base_ determines how quickly the baseline adapts to new information. By keeping this parameter low, the baseline updates at a slower pace and thus it is less prone to short-term biases.

The next neural activation pattern of each layer *n* is predicted from the distribution

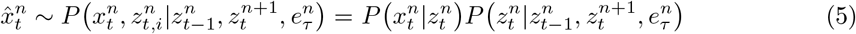

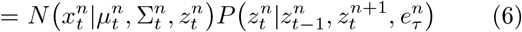

through ancestral sampling (sequential sampling of random variables each conditioned to the previous one). New feed-forward activations in layer *n* are then compared and yield the prediction error 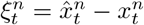. Finally, the corresponding surprise can be calculated as the negative logarithm of model evidence for this layer and this time step

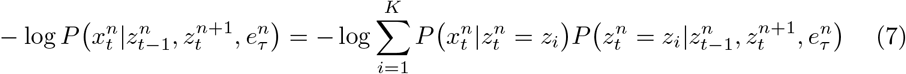

#### Splitting and merging categories

Humans are able to rearrange taxonomic relations in their memory, which often involves learning new concepts or combining two existing concepts into a single one. To capture this effect, the predict-update mechanism described so far can be also seen as a high-dimensional online clustering method based on a mixture of Gaussians. A typical approach to obtain the number of clusters in these methods is via splitting and merging existing components over time [46, 47]. Here, two components *i* and *j* at time *t* are merged iff

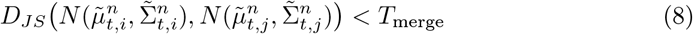

where *D_JS_* is the Jensen-Shannon divergence and *T*_merge_ > 0 is a threshold constant (hyper-parameter). In addition, the short-term state of a component *i* is split from its baseline representation if

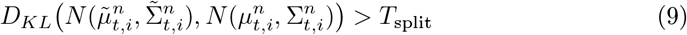

where *D_KL_* is the Kullback–Leibler divergence and *T*_split_ > *T*_merge_, with *T*_split_ another threshold constant. In this case, *D_KL_* was chosen over the symmetric *D_JS_*, as the baseline 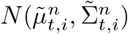 can be considered the *true* distribution of the representation of the component *i* and, thus, Eq (9) measures the amount of information lost when 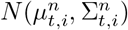 is used instead of this true baseline. In contrast, Eq (8) measures the similarity between two hitherto independent representations.

Finally, the Bayesian model described here can also be viewed as the semantic memory of an agent [48], which maintains generalized (statistical) information about different concepts in the world, including taxonomic and contiguity relations. Rather than equating semantic memory to either the generative or inference models of the brain, we view this fundamental memory system as a knowledge storage that is involved in the processing of both top-down and bottom-up signals [49] (see S1 Table and S2 Table for more details on the relation between our model and the different memory systems). The learning process followed is recursive and allows the creation of new categorical components without the need to use the same observation twice for training. However, as past observations are discarded, it lacks the ability of off-line methods to reach solutions that are optimal for all previously seen data. This issue highlights the need for the second type of memory modeled here, that can maintain approximately intact information about surprising past observations (episodes) and use this information later to improve the predictive performance of the model.

### Episodic memory formation based on perceptual surprise

Episodic memory refers to the brain’s ability to store information related to specific past experiences in the form of events and mentally relive these events either voluntarily or due to intrinsic (*free recall*) or extrinsic (*cued recall*) stimulation. Evidence suggests that the criteria used to determine which parts of the current experience are more likely to be encoded involve attention [50] and prediction error [51–53]. In the context of our model, a neural event at time *t* and layer *n* is classified as salient (or surprising) when it is very unlikely according to the current state of the system. In other words, an event is classified as salient if the generative model is not able to predict 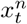 well enough. That is, iff

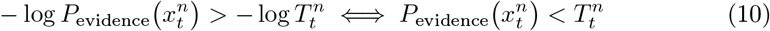

where 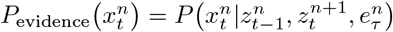 and 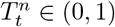 is a threshold describing the tolerated level of surprise. In a later section, we will see that this threshold can be dynamic and defines the level of attention the system pays to each hierarchical layer over time. When 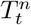 is exceeded, the tuple 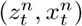 is registered as a node 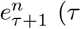 being the current number of episodic nodes in layer *n*) to a weighted, directed and acyclic graph, shown on the left-hand side of Figure 2. The value of 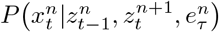 is assigned as the weight of this node, while 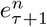 also has a direct connection to its preceding node 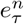 and the current node in layer *n* +1. The complete graph of episodic nodes represents the full episodic memory of the system, and provides enough information to be partially or fully retrieved in the future. The overall process of memory formation is depicted in Figure 3.

**Fig 3.**
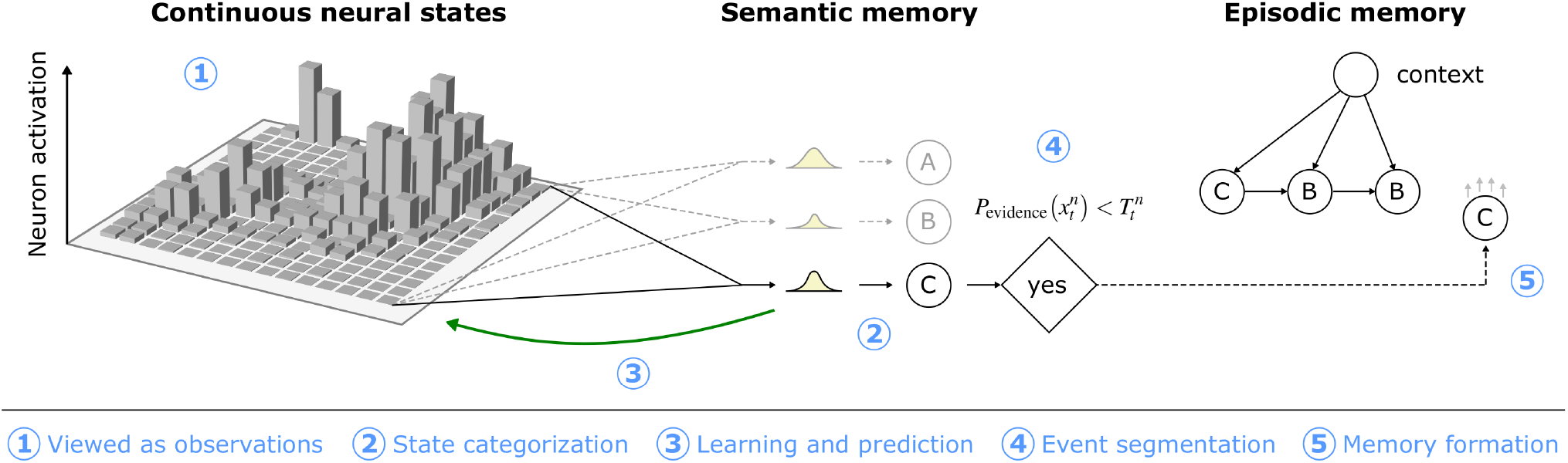
Diagram of the processes involved in episodic memory formation. At each time step, the neural state 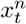 is clustered using online Gaussian mixtures and prior information. An episodic event is generated when surprise of observing 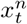 exceeds the threshold of Eq (10).

### Episodic memory recall

Information retrieval from specific episodic memories in humans can occur either spontaneously or triggered by a specific task and it is facilitated by the semantic memory [54]. For instance, when asked to report the duration of an event retrospectively, one needs to retrieve as much information related to this event as possible and fill any remaining gaps with statistical information, in order to accurately approximate the rate of perceptual change that originally occurred during this event. In this model, episodic recall is initially triggered from a single node 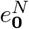 drawn from the episodic memory structure depicted on the left-hand side of Figure 2. The tree defined by all nodes connected to 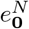 in lower layers contains the complete amount of information that has been saved for the corresponding episode. In a single recall, the process followed is summarized in Algorithm 2.

#### Algorithm 1: Tree construction during episodic memory recall

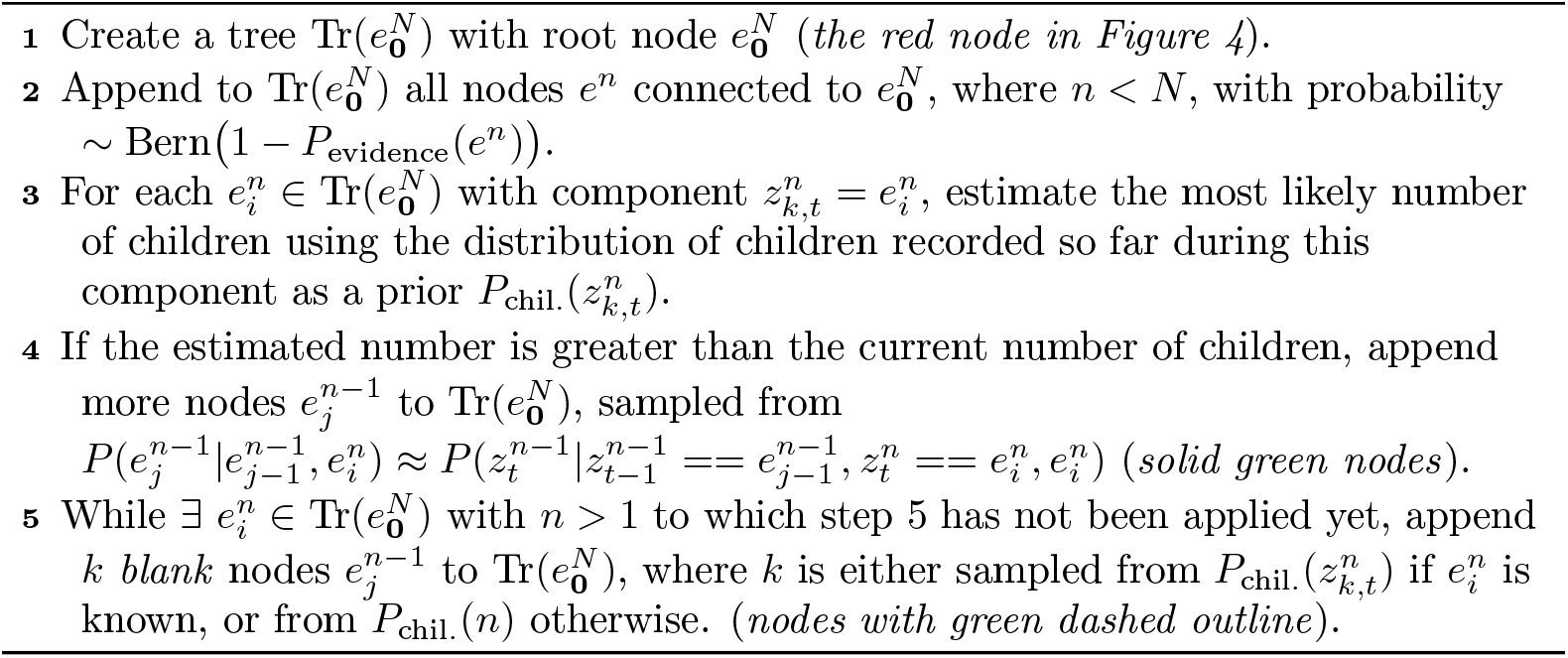

The tree 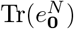 represents the salient events that occurred during 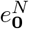 and have been currently retrieved from memory. Counting the overall nodes per layer provides an approximation of the amount of salient change in each layer during this episode, and, therefore, also an approximation of the corresponding sense of the episode’s duration. To maintain consistency in notation, let 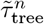 be the number of nodes in each layer *n* of the tree. Whereas the nodes added in steps 1-2 of Algorithm 2 are the only ones taken from episodic memory, alone they do not provide enough information, as they can only be fewer or equal in number to the nodes originally registered. Instead, the remaining nodes added in steps 3-5 contribute prior (semantic) information and thus fill any memory gaps.

### Estimating time

#### Prospective tasks

The inequality (10) holds mainly at times when instances of surprise occur in the system, thus detecting the presence of new, salient events. Through this mechanism, new episodic memories are formed only when their contents provide non-redundant information that could not otherwise be predicted. In [4] it was claimed that the accumulation of events of perceptual change provided an intuitive and useful proxy for perceptual surprise and belief updating and therefore could be used as the basis for human subjective time perception. As these two cognitive processes (surprise/belief updating and time perception) share the same criteria for salient event detection across time and model hierarchy, we propose that the same saliency-detection mechanism can be shared in both cases. Hence, following the current notation, the duration of an event can be estimated prospectively as a function of the number of all recorded nodes during this event, i.e.

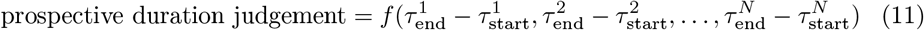

where 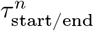 represents the number of episodic nodes in layer *n* that have been recorded at the (objective) beginning and the end of the target event respectively. After comparing different degree polynomials against human behaviour, we argue that the function *f*() can be approximated by a simple linear combination of the number of nodes (see Figure 7 and related discussion below).

#### Retrospective tasks

In order to obtain duration judgements of past events, it is necessary to recall (or at least examine) the content of the memory. Given a past event represented with a node 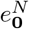, the episodic tree 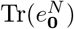 that contains both episodic and semantic (filled) information (Figure 4) represents the system’s current *best guess* of the salient events that occurred across the perceptual hierarchy during 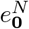, thus defining the temporal boundaries of the corresponding episode. Therefore, following the same process as for prospective duration estimates, but based on the number or *recalled* salient events in each layer for an episode, it is possible to estimate retrospective duration as:

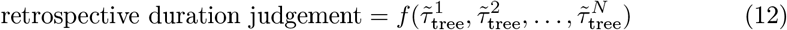

**Fig 4.**
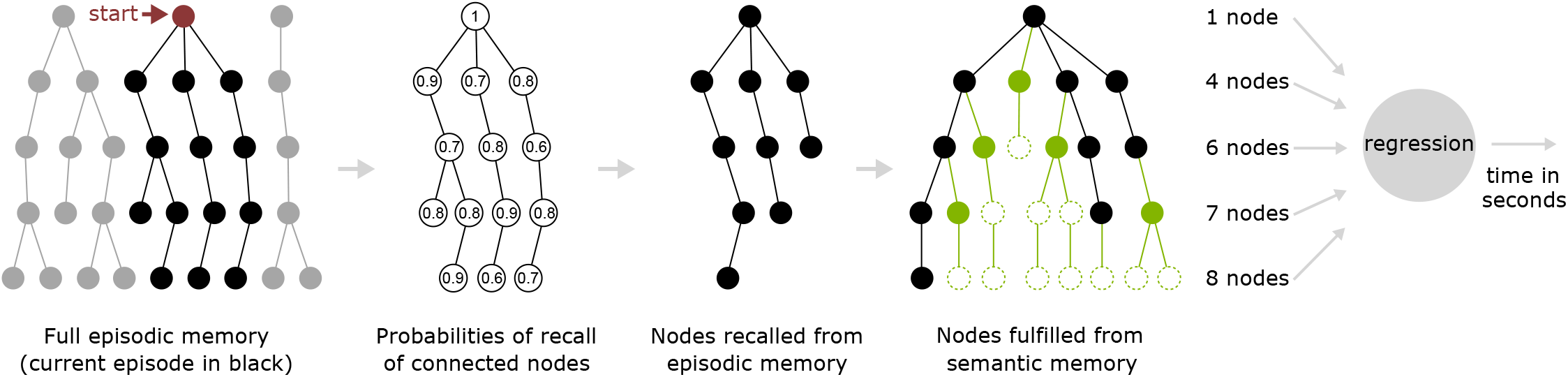
Algorithm of the steps required for episodic memory recall and retrospective duration judgements. Solid circles represent nodes with an assigned value to the categorical variable 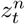 either from the episodic (black) or the semantic memory (green). Circles with the dashed outline represent nodes whose component has not been determined during recall.

In other words, the task here is to retrieve the maximum possible amount of information related to 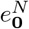 and estimate the original number of salient features during this event, in order map the latter to human-comprehensible duration units. It is worth noting that the same function *f*() can be used in both prospective and retrospective judgements as the task of mapping recorded salient events to durations remains the same in both cases.

### Attention, recall effort and cognitive load

Two factors that affect human accuracy in forming and recalling episodic memories, and subsequently performing duration judgements, are the level of attention that is paid during episode formation, and the level of effort put into recall. Intuitively, recall effort maps onto the idea that a specific portion of an episode may not be immediately recallable and describes how persistent the model should be in trying to retrieve that specific piece of memory rather than moving on without it. Here, attention and recall effort are represented through a single free parameter each, which is fitted to our experimental results.

To capture the effects of attention over time, the value 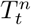 of the surprise threshold 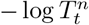 defined in Eq (10) tends to decay over time to the value 1, by obeying the equation

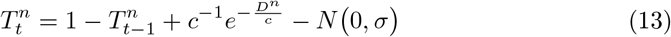

where *D^n^* indicates the number of simulation iterations since the last time the threshold in layer *n* was reset, *c* is a decaying time constant (free parameter) and *σ* indicates the level of Gaussian stochastic noise that corrupts this decay. If this threshold is exceeded, then 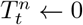 and thus 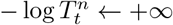. In addition, the level of recall effort (represented by the letter *ϵ*) is defined as the number of attempts to retrieve information for 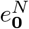, i.e. the number of times that the distribution Bern(1 − *P*_evidence_(*e^n^*)) is sampled to determine whether each node of the complete recorded episodic experience will be recalled. Our model is built on the assumption that both attention and recall effort are modulated by cognitive load. Therefore, the influence of different levels of cognitive load on duration estimation can be fully represented by different combinations of *c* and *ϵ*.

## Results

### Human experiment

We used the online platform Amazon Mechanical Turk (MTurk) to recruit 12,827 participants. Each participant watched a single video (1 - 64 seconds in duration) of a natural scene such as walking around a city, walking in the countryside or a leafy campus, or in a quiet office or a cafe (the same video stimuli were also used in [4, 7]; in these previous studies, each participant viewed a large number of videos) and reported the duration in seconds (see Figure 8). Half the participants were not informed that they would need to estimate the video duration until after they had viewed it (retrospective task group) while half were told before the trial began (prospective group). Within each group, participants completed either a low or high cognitive load trial. In the prospective group, the low-load condition required that participants estimate video duration. For the retrospective low-load condition, participants were requested to use the cues in the video to estimate what time of day the video was taking place (note that this task instruction was given prior to the video being viewed). For both groups, the high-load condition added the requirement that participants should determine whether the person recording the video was by themselves or with another person (there was never another person traveling with the person recording the video, though strangers could certainly appear).

After excluding participants we believed to be outliers (see Participants section in Methods for details), we analysed the duration judgement ratio (reported duration divided by physically presented duration) in a 2×2 ANOVA with factors of task (prospective/retrospective judgement) and cognitive load (low/high). This analysis provided evidence that the task by cognitive load interaction (*F*(1,7782) = 5.34, *p* = .02) previously reported in the literature (e.g. [10]) was present in our data. However, as some participants in this dataset reported that they were counting during estimation and we were interested in the role of external stimulation not manual counting in time perception, these participants were excluded from further analysis. The results for participants who reported they were counting (11%of participants) are presented in S2 Fig. For further details on the exclusion criteria see Methods.

Figures 5Ai-Aiv show the overall duration estimation results for each task (prospective/retrospective) and cognitive load (low/high). Our participants were capable of producing sensible duration estimates over the full 1-64 second range, with reports appearing broadly consistent with the scalar property/Weber’s law in each case (across-participant variance in estimates was roughly proportional to duration). There was some evidence of overestimation for the short intervals, compared with previous results of prospective judgements made regarding the same stimuli wherein participants completed ~ 60 trials rather than a single trial (see Figure 3a in [4]). However, there was less evidence for underestimation of longer intervals, making the present data inconsistent with Vierordt’s law/regression to the mean. The overestimation of shorter trials was likely due to the lower bound of report being limited to 0 seconds in duration, with no such corresponding upper limit for longer trials. Bounded limits in the report scale will affect duration judgements in this way [55]. Consequently, these results are consistent with the idea that Vierordt’s law/regression to the mean effects in human reports about time are the result of sequential decision processes rather than anything inherent to time perception itself [56–58].

**Fig 5.**
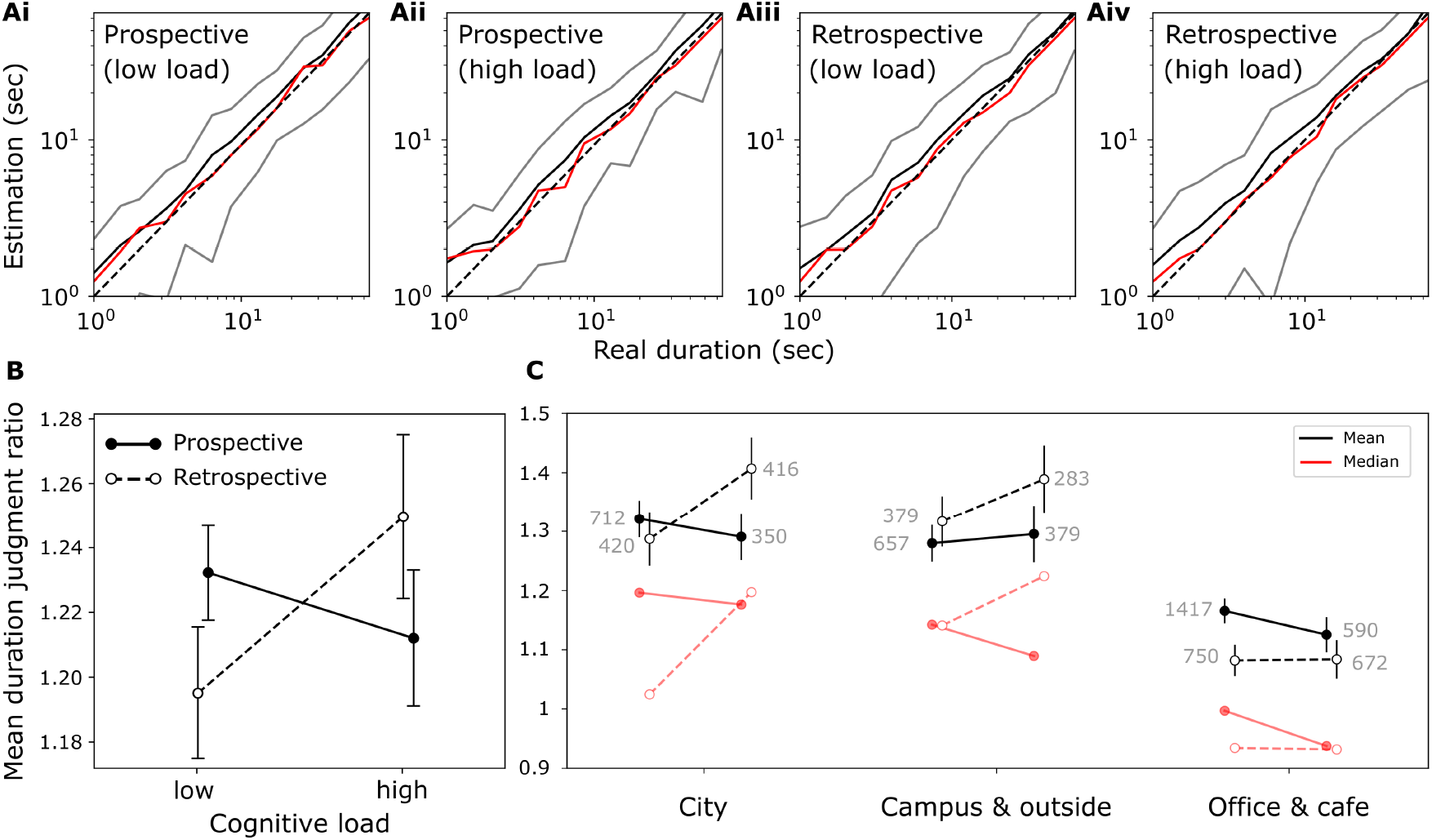
Human duration estimates according to task (pro/retrospective), cognitive load (low/high), and presented video scene (City, Campus and outside, or Office and cafe). **Ai-iv** Human duration estimation for task-cognitive load combination. The black curves represent the mean, the red is the median, and the gray is the standard deviation across all trials. **B** The mean duration judgement ratio (report versus physical duration) across all trials for each task-cognitive load combination. Broken lines/open markers indicate results from retrospective judgements, solid lines/filled markers indicate results from prospective judgements (compare with Figure 1). **C** as for **B**, but separated by scene type (City, Campus & outside, Office & cafe). The gray numbers denote the number of participants in each case, black (red) markers are means (medians).

Figure 5B shows the mean duration judgement ratio for all combinations of task (prospective/retrospective) and cognitive load (low/high). The previously reported task by cognitive load interaction [10] is still apparent in our data (compare Figure 1 and Figure 5B), though in our data it is of a smaller magnitude (after excluding counting participants; 2×2 ANOVA (*F*(1, 6973) = 3.44, *p* = .06) and is shifted towards overestimation rather than underestimation. This shift towards overestimation was likely due to our experiment using more dynamic stimuli which are known to result in longer duration estimates [4, 6, 7] (see also Figure 5C). The reason we obtain a weak task by load interaction in our data (when excluding participants who were counting) becomes clear when considering the results broken down by the different scene types that were used as stimuli (City, campus and outside, and office/cafe) (Figure 5C) - the different scene types produce qualitatively different outcomes in duration estimation. First, regardless of task and load, the degree of duration overestimation was reduced particularly for the Office and cafe scenes (for City scenes mean duration judgement ratio is 1.327 ± 0.867, for Campus & outside scenes: 1.313 ± 0.832 and for Office & cafe scenes: 1.124 ± 0.735 were the overall means and standard deviations respectively). This pattern of results was reflected in a 2×2×3 ANOVA, with task, cognitive load, and scene type as factors, which revealed a significant main effect of scene (*F*(2, 6965) = 53.07, *p* <0.001). This finding replicates previous, similar results indicating the difference in duration estimation for these naturalistic stimulus classes [4, 7, 24], but now also extends evidence for this effect to the retrospective task. Of greater interest is that the nature of the task by load interaction appears to qualitatively change with scene type. While the interaction for the City scene data (Figure 5C, *City*) was broadly similar to the overall pattern (Figure 5B), this was not the case for reports about the Office and cafe scenes. For these scenes, there was a general shift towards prospective estimates being longer than retrospective estimates, regardless of cognitive load. This pattern is the reverse of the relationship seen for reports regarding the Campus and outside scenes and was partially reflected in a significant task by scene interaction (*F*(2, 6965) = 4.10, *p* = 0.017) in the above 2×2×3 ANOVA. However, the three way (task by load by scene) interaction was not statistically significant (*F*(2, 6965) = 0.36, *p* = 0.697). Finally, the absence of a significant interaction of task and load (*F*(1, 6965) = 3.59, *p* = 0.058) when including scene as a factor reflects the similar absence in the overall analysis excluding scene as a factor, though is not interpretable beyond this superficial observation.

### The computational model replicates human behaviour

We used the exact same trials as were completed by human participants and presented in the results above as input to our model. Initially, to acquire semantic statistics of all natural scenes used, the model was exposed to 34 different 200-frame-long videos, including all scene types. Then, the same resulting semantic memory was used as the initial point for all model-based trials. The method to produce duration estimates was similar for trials from prospective and retrospective tasks (see section Estimating time). For trials completed by human participants as prospective, estimates were based on the model response to the initial input of the presented trial, by using directly the number of recorded events per layer. Trials completed by human participants as retrospective were modelled based on model-recalled episodes of the presented trial. For both cases, the difference in cognitive load was accounted for by fitting the free parameters *c* (for the effect of attention) and *ϵ* (for recall effort), described above in the section Theoretical framework, such that the relationship between model duration judgements in low and high cognitive loads follow the same slopes as the mean human duration estimates presented in Figure 5B. The parameters *c* and *ϵ* were not required to be the same for prospective and retrospective cases and, therefore, the model acted as four ‘super-participants’ each undergoing all trials of the four variants of the task. This process of fitting *c* and *ϵ* was the only part of parameter tuning in this study that was driven by human data and the resulting model behaviour that is reported in the following sections was recorded after this model fitting was performed (i.e. outside of these specific parameters, further observations of model performance are not simply a function of fine tuned fitting to human results). For further information regarding the method used to fit the free model parameters, see section Parameter tuning and S2 Appendix.

#### Duration judgements of the past and present

In the last part of model fitting, we trained a single linear regression model (using support vector regression) to map salient event accumulations to seconds on the prospective accumulation data and used it to produce time estimates for all four variants of the experiment. Figures 6Ai-iv depicts the model-produced estimates for each combination of task and cognitive load. Overall, the model was able to produce consistent estimates, with longer videos judged as longer in all cases. The variance of all reports was proportional to the reported duration, satisfying Weber’s law, while, as expected, the variance of the retrospective estimates was greater. Although overestimated, prospective model reports were highly accurate without any signs of regression to the mean. The lack of this effect, which is evident in our previous modeling approaches [4], can be attributed to the fact that the regression model here was trained under a wider range of example durations including two sets of parameters representing high and low cognitive load. Retrospective reports were also overestimated but less accurate, indicating the need for both prospective and retrospective data to be used for training the regression function *f* defined in equations (11–12) (see also below and Figure 7 for results under different training regime). Compared to human results, the variability of the model estimates was substantially lower in all cases, especially in prospective estimates.

**Fig 6.**
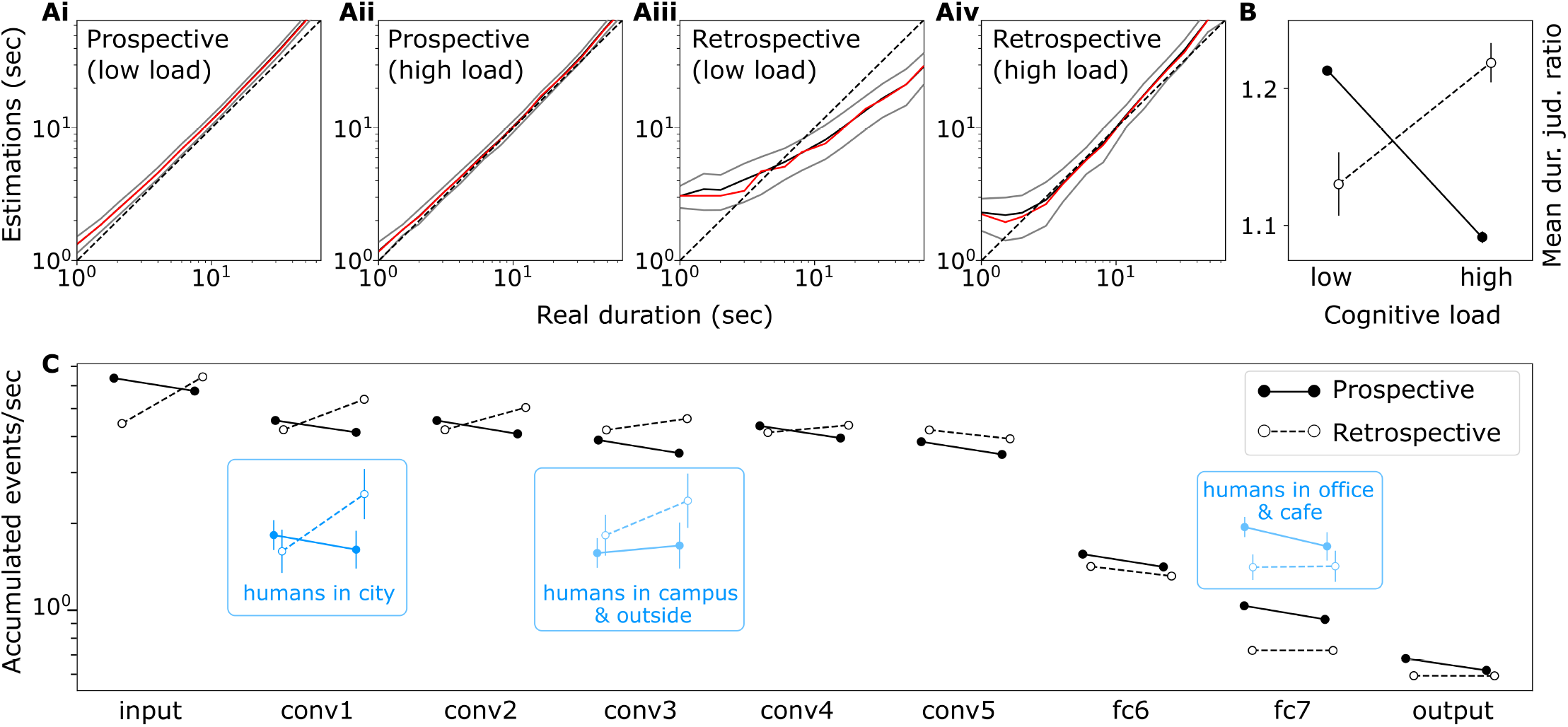
Model duration estimates in seconds or accumulated salient events by task (pro/retrospective) and cognitive load (high/low) for the same trials performed by human participants. **Ai-iv** Model duration estimates for task-cognitive load combination using a single linear regression trained only on prospective event accumulations. The black curves represent the mean, the red is the median, and the gray is the standard deviation across all trials. Estimates in seconds were obtained from a single linear model trained on objective video durations and the number of episodic nodes recorded for each trial. **B** The mean duration judgement ratio (estimate versus physical duration) over all trials for each task-cognitive load interaction, using the same linear model. Broken lines/open markers indicate results from retrospective judgements, solid lines/filled markers indicate results from prospective judgements. **C** The rate of accumulated salient events over time (per second) in the different network layers, across all scene types and separated by task and cognitive load. Note that the human data replicated from Figure 5C is in seconds and therefore is not meaningfully positioned on the y-axis - it is depicted for the purpose of comparing the task/load interaction in different scenes.

**Fig 7.**
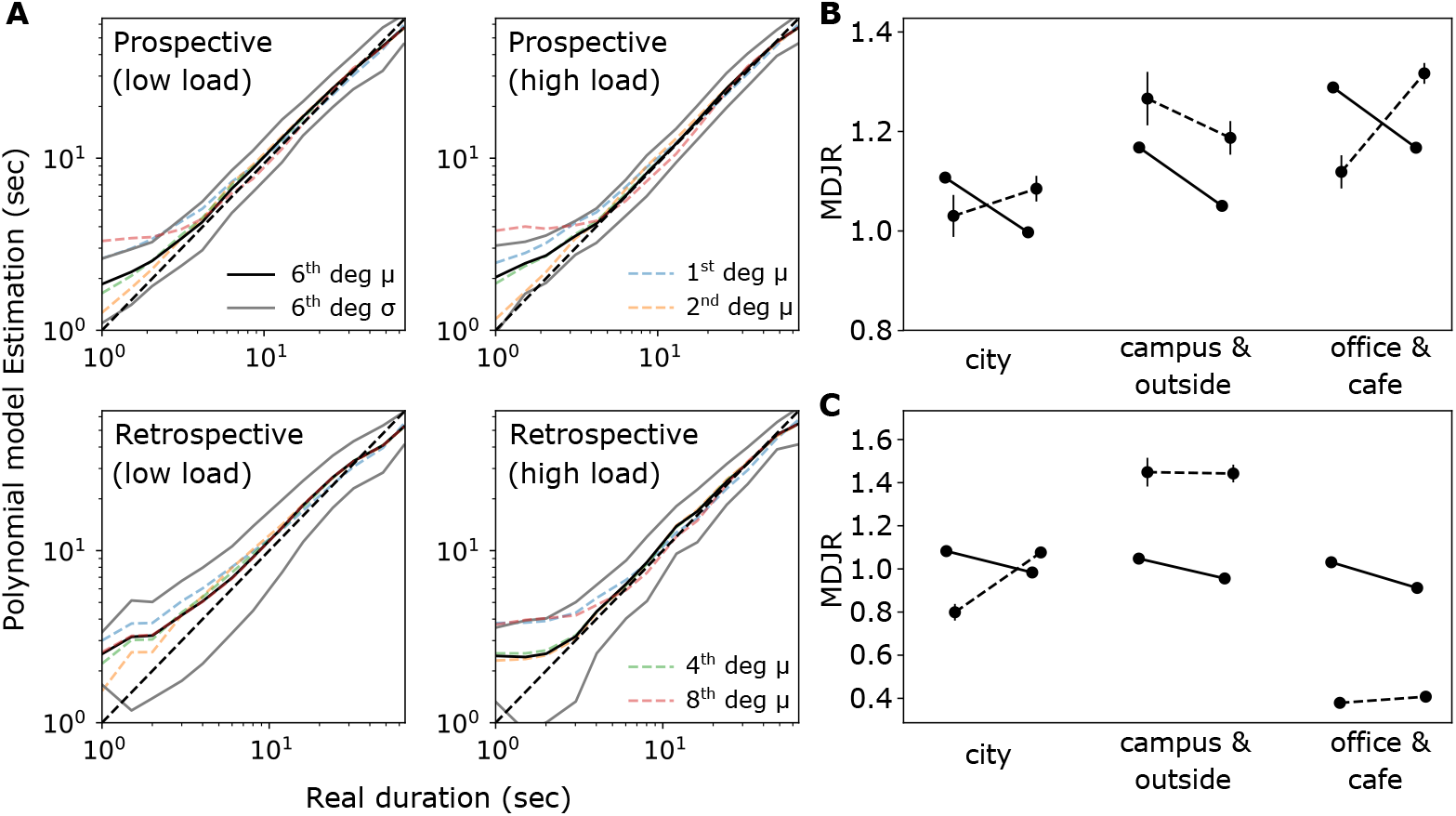
**A:** Model duration estimates for each task (pro/retrospective) and cognitive load (high/low) combination, using linear and polynomial regression trained on a single layer of salient event accumulations for different scene types, including layer *conv1, conv3* and *fc7* for trials that correspond to *city, campus & outside* and *office & cafe* respectively. In each figure panel, the solid black line shows the results for regression mapping between accumulated salient events and physical duration using a 6th degree polynomial and the grey line indicates ±1 standard deviation for estimates given by the 6th degree polynomial regression. The blue, orange, green, red broken lines indicate the results of the same regression using 1st, 2nd, 4th, and 8th degree polynomials. **B-C:** Relation between prospective and retrospective mean duration judgement ratio (MDJR) for different levels of cognitive load separated by scene type, via a single linear model using accumulated salient events in all layers (**B**) or just the layers described in Figure 6C (**C**).

Finally, the overall behaviour of the model (shown in Figure 6B) replicates the interaction of task and cognitive load demonstrated both in previous work [10] (Figure 1) and our human results in Figure 5B, with duration estimates for prospective estimates decreasing with increased cognitive load while retrospective estimates increase with increasing cognitive load.

#### Stimulus content predicts which area of the perceptual hierarchy correlates more with human duration judgments

As in previous work [4], rather than looking at the model-based estimates made following the regression of accumulated salient events into seconds, we can also inspect the accumulations directly. In our conception, these accumulations directly underlie the ‘sense of duration’ and the translation into reports in seconds is used only to enable comparison with human data. In an exploratory analysis we looked at the same breakdown by task and cognitive load as depicted in Figure 6B, but in accumulated salient events (Figure 6C) rather than in seconds, and separated by hierarchical layer in the network, and we observed that the task/load interaction varies by layer. While the clear cross-over interaction seen in Figure 6B is present in each of the lower and middle layers of the network (layers input through conv4), the interaction qualitatively changes at higher network layers (layer conv5 through output).

Most notable about these different interaction patterns is the remarkable similarities between the patterns observed in specific layers, and the patterns from human data in specific scenes. In the case of humans (Figure 5C) we see the clear cross-over task/load interaction for duration reports made about the dynamic city, while the interaction in reports made about other scenes is different. In Figure 6C we place those interaction patterns from the human results (in blue) next to the layers of the model that are qualitatively best matched. In S8 Fig we compare the cognitive load interaction between retrospective accumulated events/sec and retrospective human reports to further substantiate this match. These results suggest that duration estimation may be based on the activity in specific layers of the network for specific contexts; i.e., that humans may rely more on information captured in specific hierarchical levels of representation of an episode, given the context of this episode. For instance, stimulus representations similar to the first convolutional layer (*conv1*) of our network could be preferred for highly dynamic videos of someone walking around a busy city, representations in higher convolutional layers (such as *conv3*) in videos of medium information flow (walking around a university campus and outside), while more static videos seem to map to the last hidden layer of the network (*fc7*).

To further investigate this apparent relationship between network layer and scene type, we examined model duration estimation when accumulated salient events in only one layer of the network hierarchy were taken into account. Figure 7A shows that taking into account the single layer (identified as most similar in Figure 6C and S8 Fig); conv1 for City scenes, conv3 for Campus and outside, fc7 for Office and cafe scenes), estimation performance increases and becomes more similar to the human results than that shown in Figure 6A wherein the regression was fitted on all accumulation in all network layers. For a comparison between the two computational models in the same axis as the human data and the previous model in [4] see S7 Fig.

Model comparison using the Akaike information criterion (AIC) showed that the best performing model (both in terms of capturing human behaviour and estimating physical duration) trained to map between the accumulated salient events in a specific layer and physical duration using a 6th degree polynomial (solid black line in Figure 7A); however, note that even a linear model (broken blue line) performs well. Moreover, the interaction between task and cognitive load by scene is much more consistent with that seen in the human results (Figure 5C) when the regression (linear in this case) is trained only on the qualitatively matched single layer accumulation (Figure 7C) compared to when it is trained on accumulation in all layers (Figure 7B). All AIC scores for the different candidate models compared to both human reports and physical durations are reported in S3 Table.

Overall, these results show that it is possible to fit a simple linear model, using only a single layer per scene type as input (only two parameters x 3 scene types = 6 parameters in total) and real durations as the target (i.e. without using human data in fitting), that can replicate the complex pattern of interactions between task, cognitive load, and scene-type that we found in human duration estimates (Figure 5C) for these same naturalistic video inputs. Assuming such a simple scenario, an interesting direction of future research then becomes the questions of *how* (and *why*) a single appropriate layer is selected. In the Supporting Information we explore several possibilities for layer selection based on network behaviour in different layers in different contexts (i.e. scene types). These approaches include variability of accumulation in a layer, uniqueness of components for a given memory episode across layer, and accuracy of recall across layer (S5 Fig). All three approaches exhibit differences across network layer and therefore may provide potential foundations for a decision process that selects the specific network layer from which to estimate duration, given a specific context. Further research on this topic is required, while, due to the post-hoc nature of the results in this section, our conclusions should be further validated by another study with a different set of videos that replicate our findings.

#### The role of attention and recall effort

In [4], it was shown that attention *c* has an important effect when modeling prospective duration judgements. This relation is verified again here through the behaviour of the current computational model, as *c* was the only free parameter used to model differences in cognitive load. However, in retrospective judgements parameter tuning of *c* had minimal effect and was inadequate to produce model reports consistent with the human biases due to the cognitive load differences (see S6 Fig, especially *B*). Therefore, we kept the *c* fixed for both retrospective cases of cognitive load (values of all parameters are reported in Methods).

In contrast, our analysis indicates that the parameter *ϵ*, used to control the level of recall effort, has a substantial impact on the hierarchical structure of the memories being recalled and can be tuned to model human cognitive load biases (see S1 Appendix). It predicts that committing more effort to recalling an episode should generally lead to greater overestimation of the episode duration and that visual episodes which are rich in content should be more overestimated. Our current experiments however do not provide substantial evidence of a direct relation between high cognitive load and high recall effort and further work is required to study this relationship.

## Discussion

The primary contribution of this work is a novel framework (and a computational implementation) for viewing human time perception via episodic memory in both prospective and retrospective cases. First, we designed a neural architecture that couples predictive processing, time perception, episodic and short-term memory. Additionally, we ran a large-scale human experiment where we recorded duration judgements from ~ 13, 000 participants, under two manipulations including both prospective and retrospective timing. Using the proposed model, we were able to reproduce the characteristic patterns of human duration judgement biases (estimated from ~ 7000 of the human participants after excluding outliers) in all experimental manipulations, taking as input only the rate of salient events detected within a hierarchical perceptual processing network. Hence, we showed that using a single computational process to estimate duration both during ongoing perceptual input (prospective) and based on recalled memory of events (retrospective), our model reconciles divergent approaches and findings in the time perception literature and accounts for observed differences in human duration perception across tasks, cognitive load, attention, and stimulus content.

A number of conceptual similarities connect the current proposal and the computational model introduced by Roseboom et al. [4]. Both studies are based on the fundamental proposal that time can be estimated in tracking activity in perceptual modalities [7, 24], they use the same type of natural stimulus, and the same mechanism for detecting salient events. Although in the previous model surprise was modelled by simple Euclidean distance between neuron activation in consecutive frames, here it is defined in a Bayesian fashion, as the negative log-model evidence. Crucially, the study in [4] did not account for the concepts of episodic, semantic or short-term memories and, therefore, was incapable of 1) reproducing results for tasks that rely on memory, i.e. retrospective tasks, and 2) viewing accumulation of salient changes for time estimation as fundamentally linked to event segmentation for memory formation. Finally, the current human experiment was designed for one trial per participant (as required to keep participants naive about the question of time for retrospective judgements), contained a manipulation of cognitive load, and checked for and excluded from analysis where necessary participants who reported that they were counting time. All these design differences provided for additional insights not possible based on the human data of previous studies, such as the lack of the characteristic regression to the mean in duration estimates (Vierordt’s law).

### Deep predictive processing with episodic memory

The literature contains various neural implementations of predictive processing, including deep generative models [59–61]. and models that aim to replace backpropagation with local learning rules [62, 63]. There are, however, a number of significant differences between existing computational models and the approach proposed here that are necessary for modelling retrospective timing tasks. Most importantly, the present model incorporates predictive coding with episodic memory (see Figure 2). Local prior predictions of the latent states 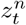, which are part of the prediction-update loops in each hierarchical layer are jointly conditioned on episodic, short-term semantic and long-term semantic information, as shown in equations (5–6) and (4). Furthermore, the present model incorporates a semantic mechanism that categorizes continuous states into discrete indicative variables. These indicative variables can be viewed as “labels” of the continuous representations and recorded events. We argue that the processes of event segmentation and state categorization are fundamental for the formation of new episodic memories (see Figure 3 and next sections in the Discussion). Finally, the model presented here is Bayesian, in the sense that it is realized via a set of random variables, whose corresponding probability distributions are estimated via approximate Bayesian inference.

### The subjective timescale of time perception and episodic memory

Our model shows that the dynamics of human perception could update over two distinct timescales; short-term and semantic memory systems follow the rate of changes in the real world (right-hand side of Figure 2), whereas the episodic memory system follows the *subjective* rate of captured salient events in a given hierarchical representation (left-hand side of Figure 2). We argue that human perception of duration (in all cases) resides entirely in the latter, hidden temporal structure and, therefore, has no direct connection to the physical passage of time [64]. This approach follows from the intuitive basis that human perception of time is characterised more by its common deviations from veridicality (see [24]), rather than acting like a clock that attempts to track elapsing physical time (e.g. [13, 65]). Without a direct connection to physical time, human tasks that involve subjective duration reports must rely either on episodic memory formation (prospective time) or recall (retrospective time) processes. To communicate estimates of duration or relate subjective time to the physical world (like timing how long it may take to walk to your nearby bus stop), the brain need only employ a read-out network that learns to map the length of a memory to a standard unit, such as seconds. The emergent human-like performance in Figure 6B and Figure 7 using a linear model as the function *f*(.) of equations (11–12) supports this view and highlights the potential simplicity of this read-out process, which could be performed by even single task-, content- and unit-specific neuronal cells. Such a read-out process would have a wide repertoire of available and partially overlapping resources across the perceptual hierarchy to potentially use. Therefore, it is reasonable to speculate that individual hierarchical layers might be employed, since this would require less computational resources. These layers could be different for different stimulus content, as in the case of our model shown in Figure 7, if we assume that different levels of perceptual representations might be more informative for different stimulus content. In busy city scenes for instance, where more high-layer events are recorded than on average, relying on a more consistent low perceptual layer could lead to a lower uncertainty in estimations.

It is worth stressing the difference between our proposal and proposals of mechanisms that involve the direct (or indirect) tracking of physical time. As with any other dimensions of experience, the human brain constantly receives redundant time-related information. For example, photoreceptor cells in the retina are driven by light, and low-level neural dynamics are governed by physical time as neurons are bound by electrophysiological constraints. However, most of the information received by the retina is lost in the pathways of the visual cortex; according to predictive processing, sensory information is propagated only or largely in the case that it cannot be predicted by the brain’s model of the hidden causes of sensory input [25]. Contemporary interpretations show that this mechanism optimizes energy efficiency in terms of information processing and storage [35, 66, 67], while it has been argued that it even constitutes a fundamental requirement of (biological) self-organized systems to achieve homeostatic control, or life [68–70]. Here we view time as simply one more source of observations that the brain needs to reconcile with its internal model of the world. Indeed, our proposal takes time perception one step further from the physical world than other sensory modalities, as the cognitive architecture described in Figure 2 and S1 Fig does not require an explicit representation of time, or indeed any ‘time sensors’ (compare, e.g., vision). Instead, through the processes of event segmentation and episodic memory management, salient temporal information is encoded in a parsimonious fashion and remains available for most time-related tasks. Exceptions to this basis for temporal processing might include pattern timing tasks for short intervals, where neural network states, analogous to photoreceptor cells for vision, can be more useful [71, 72].

Finally, our model attempts to reconstruct previous stimulation based on a combination of what can be recalled from a given previous episode and the properties (in terms of stimulation) of similar episodes (referred to as semantic memory). However, neither the formation nor recall processes contain the contribution of any explicit knowledge of previous duration, such as the explicit knowledge that a previously experienced episode lasted 45 seconds. While humans have the ability to retain and recall such information, potentially in isolation of the content that may have originally underpinned the time estimation [73], it is beyond the scope of our present modelling work to incorporate such influences. While of great interest, such work would require a much deeper understanding of the interaction of experience, language, and memory than we currently possess, though this is likely not outside the scope of an extended predictive processing approach [74].

### Time perception and episodic event segmentation

Perceived time in our model is determined by the frequency of occurrence of salient events over network layers for an epoch. These events can be interpreted as episode boundaries (at each layer) – the changes in content, with the relevant content type defined relative to network layer (more complex content at higher layers). In this way, our model provides a method for event segmentation in defining memory episode extent.

The relationship between time perception and memory segmentation goes back at least decades (e.g. [75–78]), though a formal description of event segmentation in episodic memory has only taken a prominent position more recently (e.g. event segmentation theory [79, 80]). Extensive work in both humans [81–85] and animals [86, 87] has identified (changes in) hippocampal activity as related to human reported (annotated) event boundaries and, more precisely, demonstrated the key role that (lateral) entorhinal cortex plays in the temporal arrangement of events for episodic memory [84, 85, 87]. It has been suggested that activity reflecting event segmentation is based on prediction error [88–91], placing the determination of event segmentation firmly within a predictive processing framework, in line with that demonstrated here.

Predictive processing has the distinct advantage over alternative modelling approaches that a definition of salient event (or event boundary) is possible without involving controversies about whether the processes that define time perception and episodic memory formation are related more to dynamics (or intensity) of external stimulation, or to internal states such as attention or task-specific context [8, 92] - instead, salience is always defined as a continual trade-off between stimulation and expectation. Consequently, while our model accommodates the powerful and well-established intuition that change in perceptual stimulation underlies subjective time [1, 2, 4], the context of stimulation and prior knowledge of similar scenarios are also fundamental for determining what constitutes *salience* (see also section Predictive processing neural architecture for perception). In addition, this formulation allows the current model to form salient events and, thus, produce prospective time estimates of an episode, even in cases that either noise or a static image is presented to the model across the entire episode duration. This is possible due to the decaying threshold of surprise in Eq (13) that captures the effects of attention in memory formation. In more complex implementations of the current model, this threshold is expected to play less significant role, as surprise across the perceptual hierarchy could be driven by more complex intrinsic processes, such as memory consolidation or mental time travel.

Finally, although the current model relies on the existence of event boundaries in continuous experience, it is worth noting that the thresholding mechanism described in Eq (10) and Figure 3 (step 4) could be replaced by a mechansim that regulates the originally recorded perceptual surprise 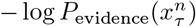 based on the recency of the event (*t* − *τ*). Without an attention threshold, all instances of a given stimulus would be initially recorded in episodic memory, along with an initial memory weight. This memory weight could be further scaled to simulate the effect of attention. As the recorded instances become less recent, however, the memory weight of instances with low initial surprise will be reduced to zero creating boundaries between areas where the initially surprise was significantly higher, i.e. forming events. Hence, except for being significantly more computationally expensive, this approach has practically the same effect as temporal discretization with 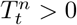.

### Hierarchy of temporal processing

Previous studies have highlighted the hierarchical nature of temporal processing in the brain, with the timescale of processing at lower cortical levels (e.g. primary sensory cortices) being shorter and more stereotyped than processing at higher cortical areas (e.g. superior temporal sulcus and frontal eye field [82, 93, 94]). Beyond simply hierarchy alone, episodic memory is nested with more complex events comprised of smaller-scale and less complex events that provide internal structure to events as defined at the highest level (*partonomic* hierarchy [91]). The tree-based structure (see Figure 4) of our model is nested in its configuration and the perceptual classification network at the core of the network exhibits a hierarchical distribution of representational complexity (nodes in lower layers are selectively responsive to basic image features like edges, while nodes in higher layers are responsive to complex whole-object like features) – a feature shared with biological perceptual processing [16, 18]. The depth and distribution of representational complexity across our model hierarchy allows us to reproduce the differences in human duration report in different contexts – dynamic city scenes versus relatively static office scenes. Furthermore, we demonstrate that time perception may be accomplished by accumulating salient events in only some portion of the neural hierarchy, relevant to the ongoing context at the time (Figures 6C and 7B and C). This possibility is consistent with previous findings that the temporal dynamics of salient event transitions can be detected within the region of the brain relevant to the temporal scale of that event. For example, neural dynamics related to event boundaries in basic stimulus properties such as motion are detectable in lower level sensory areas like human MT [81, 82, 95] while higher complexity events will be detectable in the dynamics of higher cortical regions [82]. A context-related cortical specificity for salient events is also consistent with the finding that only the accumulation of salient events within the relevant sensory areas (ventral visual areas when watching silent videos) can reproduce human reports of subjective duration [24].

While structurally equipped to produce many properties of episodic memory, here we have only tested the time perception abilities of our model. Further work is required to determine the depth to which our model (or our modelling approach more generally) reproduces the many other known features of human episodic memory (see [91]). In particular, future work will need to address the precise degree to which important concepts in the memory literature such as primacy, and recency and contiguity [96–98] are produced in our memory model. Though it is worth noting that contiguity and forward asymmetry are preserved in principle by the model construction, with related events recorded and reconstructed in sequence within hierarchically nested episodes (see Figure 4). Further work is needed to fully relate these features of our model to findings from the literature that rely on word list tasks or similar rather than naturalistic visual sequences.

### Interval timing via non-temporal Bayesian inference

Our process of episodic recall recursively estimates the number of children nodes for each recalled event (see step 3 of Algorithm 2). This process can be instantiated as a form of Bayesian inference over the duration of events, wherein new observations of the number of nodes for which an event can be parent updates the model’s prior knowledge. It is worth noting that the mechanism of prospective timing proposed here is also consistent with this interpretation and could support inference, if prior semantic knowledge is combined with the current number of recorded (or recalled) events. Although the simulation results of the prospective task presented here are based only on current accumulations, as prior knowledge would have a negligible effect on single-trial experiments, this recursive inference process could potentially account for other known effects in the time perception literature, such as effects generally seen as regression to the mean of estimation [56–58, 99]. In addition, the Bayesian treatment of accumulated events allows for more sources of information to be integrated during estimations. This may be particularly important for very small intervals where more rhythmic mechanisms might be expected to provide meaningful information for duration related tasks [72]. This proposal might be evidenced in various simple ways, such as finding differences in Vierordt’s law/regression to the mean in a retrospective timing task, when episodic memory recall is manipulated. Finally, the accumulation of salient events defined by the thresholding mechanism in Eq (10) can be seen as an approximation of the amount by which the system’s prior beliefs *P*(*z^n^, μ^n^*, Σ^*n*^) update under a specific period of time. This view allows us to define an analytical form of perceptual change using tools from information geometry, such as the Fisher’s information metric between belief states [100], and connects the fundamental notion of information to human perception of time, under the Bayesian framework.

### Further study required

Like any mathematical model of a biological process, the present implementation is accompanied by a number of limitations. Since raw neural activation 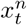, and not prediction errors 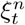, is what propagates to higher layers in the network, the system can maintain the effect of a new sensory event for an unrealistic amount of time. Hence, the dynamical behaviour of the neural activity here should not be confused with biological neural activity, a fact that precludes predictions at that low level of description. This issue could potentially be counterbalanced by the existence of short-term plasticity in equations (2–4) that quickly reduces prediction errors, if 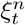 is thought to represent neural activation. However, more investigation is required to assess this relation.

In addition, the network presented here is based only on visual processing. It does not currently have the ability to integrate information from different sensory modalities, to parse higher level visual information (such as the type of room of the current environment), to perform basic spatial navigation, nor to pay attention to only a specific part of a given stimulus. Nevertheless, given the nature of our predictive processing approach, and previous findings demonstrating the similar structure of neural activity related to event transitions in different sensory modalities (combinations of modalities) [82], we anticipate that our approach is readily extendable to incorporate these additional features under a broader active inference approach [36].

Finally, our human ability to reconstruct past experiences is considered closely intertwined to our ability to create completely fictitious scenarios, which either could have happened, or might happen in the future [101, 102]. These two abilities are often referred to together as mental time travel [103]. Research in this field indicates a strong connection between recalling past experiences, semantic memory and imagining the outcomes in potential future scenarios [104, 105], an undoubtedly crucial aspect of our decision making [36, 106]. The mechanism for episodic memory recall presented here can readily encompass the notion of imagination, if the selected root node of the tree structure *Tr*(.) is not taken from existing episodic memory, but it is simply filled by a component 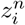, as in the case of the *blank* nodes depicted in Figure 4. Under this small modification, the recall mechanism would trigger the generation of a tree where all remaining nodes are fulfilled based on prior (semantic) beliefs. Thus our model presents a platform with the necessary components to investigate a wide range of theories related to memory and time such as imagination [107], dreaming [108], and hallucinations [109], based only on such variations of how the episodic tree is constructed or recalled. Such a platform would likely prove highly valuable for both psychological and neuroscientific work on these topics, but also for the expanding fields of deep neural networks and reinforcement learning [110].

## Methods

### Human experiment

#### Experimental procedure

The human experiment was conducted using the online platform Amazon Mechanical Turk (MTurk). The participants were asked to watch a single video using the full size of their home computer monitor and then answer a series of questions including a report of the perceived duration of the video. Each participant completed a single trial that lasted from 1 to 64 seconds. Any visual information regarding the real duration of the video was removed from all trials, apart from a description of the video using the expression ‘very short’, applied to all video durations. 50%of the participants were not instructed that the experiment involved duration judgements before the final questionnaire (see the description of the tasks bellow).

#### The platform

The large number of participants required for this experiment (one participant per trial) makes it impracticable for a typical lab setting. Amazon’s Mturk platform provides a satisfactory alternative approach [111]. It maintains a large and diverse worldwide pool of about 500,000 potential subjects, with a large portion of this pool shown to be a good representation of the general population of the United States [112].

#### The tasks

The task comprised a single trial with four different variations, one of which was randomly assigned to each participant. These task variants were designed to allow for two different levels of cognitive load, as well as prospective and retrospective duration judgements. Trials comprised three successive screens. In screen 1, participants were told they will see a very short video and then, they will be asked a number of questions regarding the instructions given in Figure 8. In screen 2, the selected video was automatically played without any controls or further instructions and, once finished, the screen 3 spawned automatically with the corresponding questionnaire.

**Fig 8.**
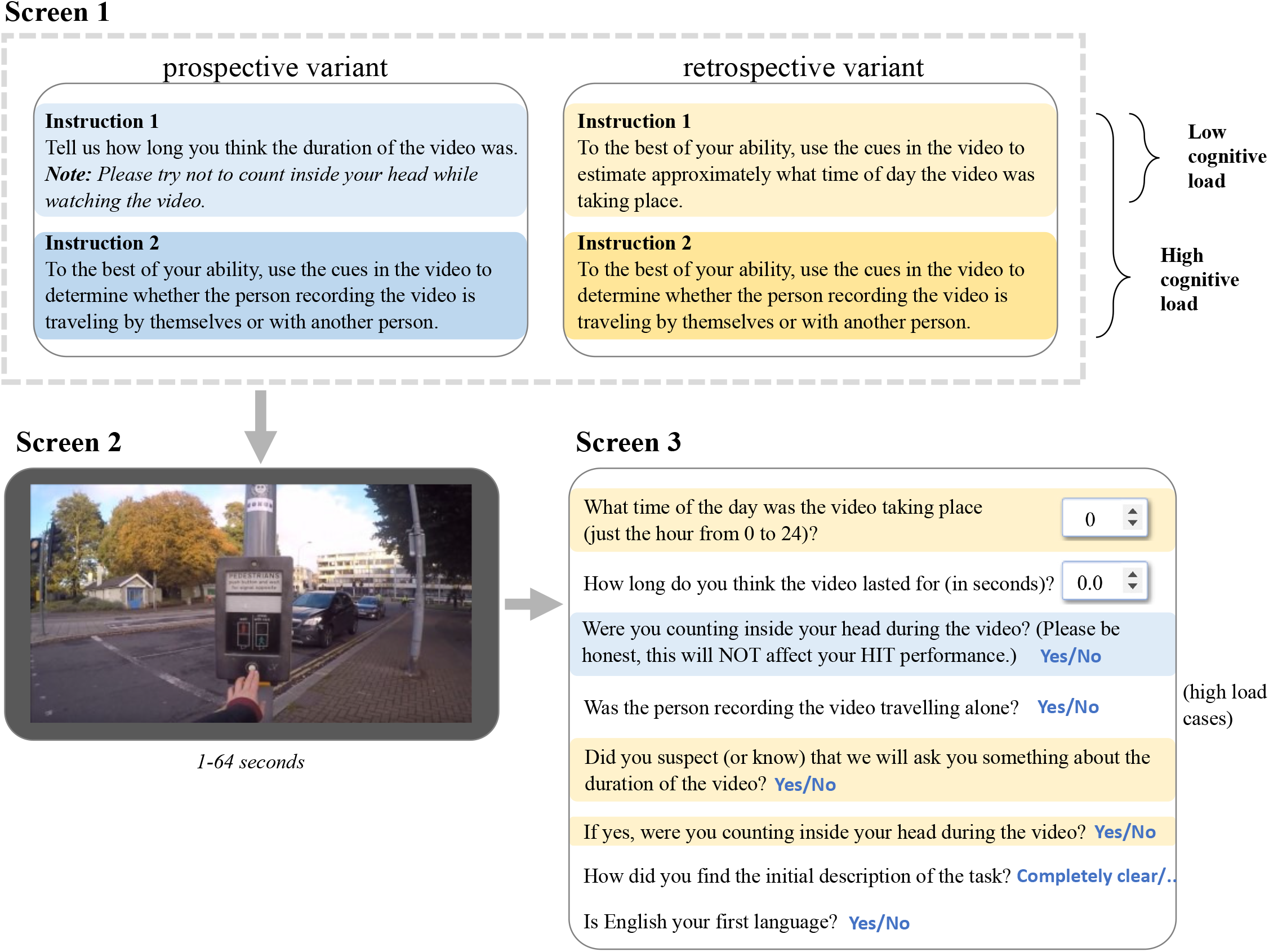
The main steps of the trials including instructions the video and the final questionnaire, for the four variants of the task.

#### Participants

Using the MTurk platform, we recorded data of 12, 827 unique individuals who performed one of the four variations of the experiment. Subjects were based in the United States, Canada, Australia, New Zealand, the EU and India and they received payment of US$0.25 per trial. Subjects were anonymous, over 18 years of age, they reported to have normal or corrected-to-normal eyesight and they provided their consent by agreeing in a web form to the terms and conditions approved by the University of Sussex’s research ethics committee (C-REC; approval code ER/WJR22/9).

Out of the total number of participants, the primary results presented here excluded subjects who (1) were in the retrospective condition but reported that they realized during the trial that they would need to respond about duration, or (2) reported after the trial that they found the experiment instructions either ‘completely unclear’ or ‘somewhat unclear’. To reduce the effect of the uncontrollable factors in the environment of the experiment, participants were also removed if (3) the video playback was different from the desired real duration for that trial, due to slow internet connectivity or any other factors. In particular, each trial was rejected if either

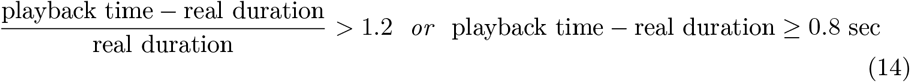

Furthermore, although participants of the prospective variant of the experiment were instructed to try not to count in seconds during the trial, when asked in the final questionnaire some reported they could not resist counting. Counting was also reported by a small group of participants of the retrospective variant, who suspected they would be asked about the duration of the video. All these participants were also removed from the main sample.

After the exclusions (1-3), the sample comprised 8072 participants and, after removing 933 who reporting counting, the remaining sample comprised 7139 participants. Removing outliers for the reported durations (any duration such that report 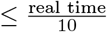 or report ≥ 10 × real time) left the final sample used for the results throughout this manuscript, which comprised *n* = 6, 977 non-counting participants (and trials) or *n* = 7, 786 when counting participants are included. Outliers were also removed from the group of counting participants (*n* = 809) in order to produce S2 Fig. Outliers and counting participants were also removed from the four distinct groups illustrated in S3 Fig that compare the behaviour of participants that reported different level of understanding of the initial task instructions. Finally, we also recorded which participants were native English speakers, as well as the screen and web browser sizes, to ensure the trials were run in a wide field of view. An overview of the demographics of the final sample can be seen in S4 Fig. The resulting dataset can be found in the repository^2^.

### Computational model implementation

#### Computational model implementation

To implement the inference model 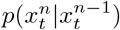, as depicted in S1 Fig, we used a well-studied convolutional network defined by Krizhevsky and colleagues [19]. Deep convolutional networks have hierarchical structure inspired by and broadly resembling the human/primate visual cortex [16–18, 113], and have been shown to perform very well in visual classification tasks [19, 20]. This network was pre-trained to classify natural images into 1000 different categories of objects and animals [114]. It comprises 9 neuron layers of which 5 hidden layers are connected with convolutions (*conv1-5*) and 3 layers with all-to-all connections (*fc6-7* and *output*).

To predict the next activation pattern of each layer *n*, the method of ancestral sampling was used. Instead of sampling from the joint distribution described in Eq 5 directly, the algorithm initially samples from 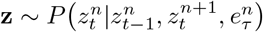. As the variable 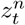 is discrete, this distribution is approximated by a table that tracks the number of transitions to different values of 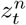 over time. Then, **z** is used to condition 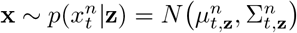. To approximate the model evidence for each Gaussian component, needed for Eq (7), the multivariate Gaussian probability density function is used instead. Finally, to approximate the initial covariance matrix 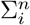 for each new component *i*, the overall covariance 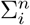 for the layer *n* is tracked and used.

#### Parameter tuning

The hyper-parameters related to the generative model (Table 1) were obtained via optimization with the double objective of (1) reducing the average surprise 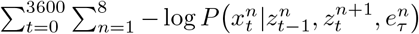 and (2) maximizing clustering accuracy in the components of the output layer *z*^8^. As a dataset, a 120-second video in an office environment was manually labeled, in order to capture the most prominent object per frame. The optimization method used was the multi-objective Covariance Matrix Adaptation Evolution Strategy (MO-CMA-ES) [115] a widely-used derivative-free algorithm, ideal for non-linear and non-convex parameter optimization. The parameters for the attention and recall mechanisms were also tuned using CMA-ES, following the principles of *c*_low prosp_ < *c*_high prosp_ < *c*_retrospective_ and *ϵ*_low load_ *ϵ*_high load_, such that the overall relation between tasks and cognitive load levels, shown in Figure 5B, is approximated. These constraints represent the working assumptions that (1) less attention to the main task is expected in cases that humans are distracted with multiple tasks and that (2) human subjects are not expected to be more persistent in trying to retrieve a specific piece of memory, in cases that they were less distracted during the original experience. The complete model training process is described in pseudo-code in S2 Appendix.

**Table 1.**
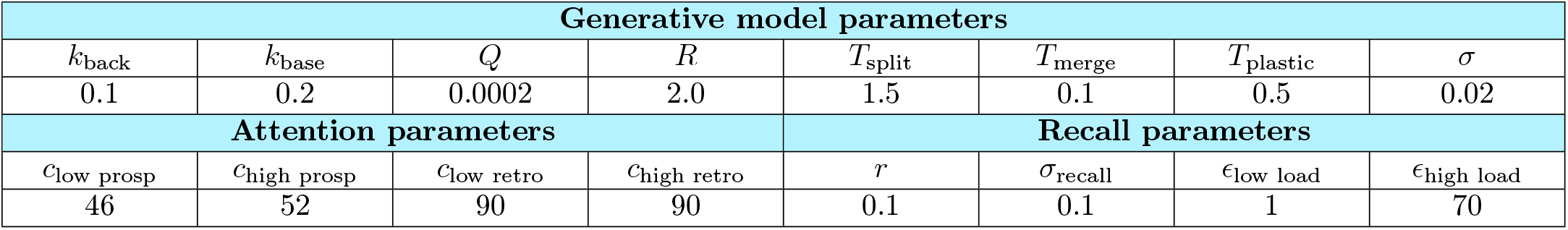
Model hyper-parameters

| Generative model parameters |  |  |  |  |  |  |  |
| --- | --- | --- | --- | --- | --- | --- | --- |
| $k_{\text{back}}$ | $k_{\text{base}}$ | $Q$ | $R$ | $T_{\text{split}}$ | $T_{\text{merge}}$ | $T_{\text{plastic}}$ | $\sigma$ |
| 0.1 | 0.2 | 0.0002 | 2.0 | 1.5 | 0.1 | 0.5 | 0.02 |
| Attention parameters |  |  |  | Recall parameters |  |  |  |
| $c_{\text{low prosp}}$ | $c_{\text{high prosp}}$ | $c_{\text{low retro}}$ | $c_{\text{high retro}}$ | $r$ | $\sigma_{\text{recall}}$ | $\epsilon_{\text{low load}}$ | $\epsilon_{\text{high load}}$ |
| 46 | 52 | 90 | 90 | 0.1 | 0.1 | 1 | 70 |

<sup>2</sup>Human dataset: https://github.com/zfountas/prospective-retrospective-data

During the training process described in S2 Appendix, the accumulated events for the four variants of the experiment, as well as regression weights for the model presented in Figure 6 were recorded. these accumulated events were later also used to fit the single-layer polynomial models presented in Figure 7. To train the single-layer models, the dataset of event accumulations was divided into 4 independent sub-sets, one per each variant of the experiment and the function *f* defined in equations (11–12) was fitted 4 times with all data of each 4 sub-sets, using again the real durations of the video and ordinary least-squares polynomial regression. As in the multi-layer regression model, the resulting 4 models were tested in human reports of the four experiment variants respectively.

Finally, the programming language used throughout the modeling implementation is Python 3.7 and the machine learning library TensorFlow 1.14. The source code can be found on the repository^3^. The analysis of the results was also performed in Python 3.7. The scripts used for the analysis of the human data can be found on the repository that contains the human dataset, while the scripts that perform analysis of the model behaviour, along with the resulting dataset of the computational model are included in the repository of the model’s source code.

## Supporting information

**S1 Appendix. Analysis of the role of effort in memory recall.**

**S2 Appendix. Model training process.**

**S1 Fig.**
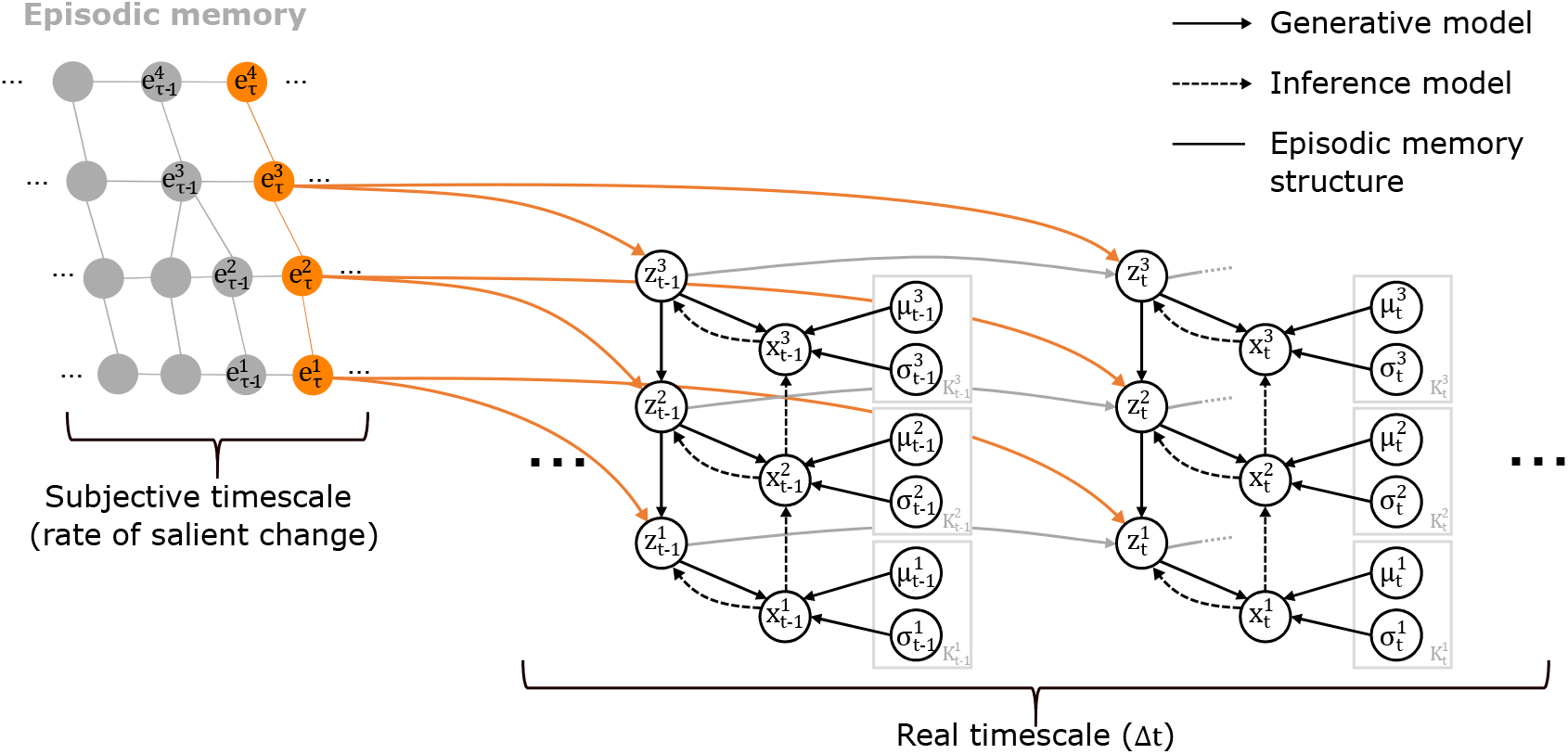
Full probabilistic graphical representation of the predictive processing model p. The random variables 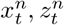 correspond to different hierarchical representations of the input, continuous and categorical respectively, the variables 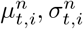 are the parameters of the Gaussian component *i* and the variables 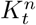 represent the number of components at time *t*. All solid-line arrows represent conditional dependencies: black arrows denote dependencies between random variables within the same simulation time step, red arrows highlight dependencies between variables with distributions that evolve over different time scales and gray arrows denote dependencies across different time steps.

**S2 Fig.**
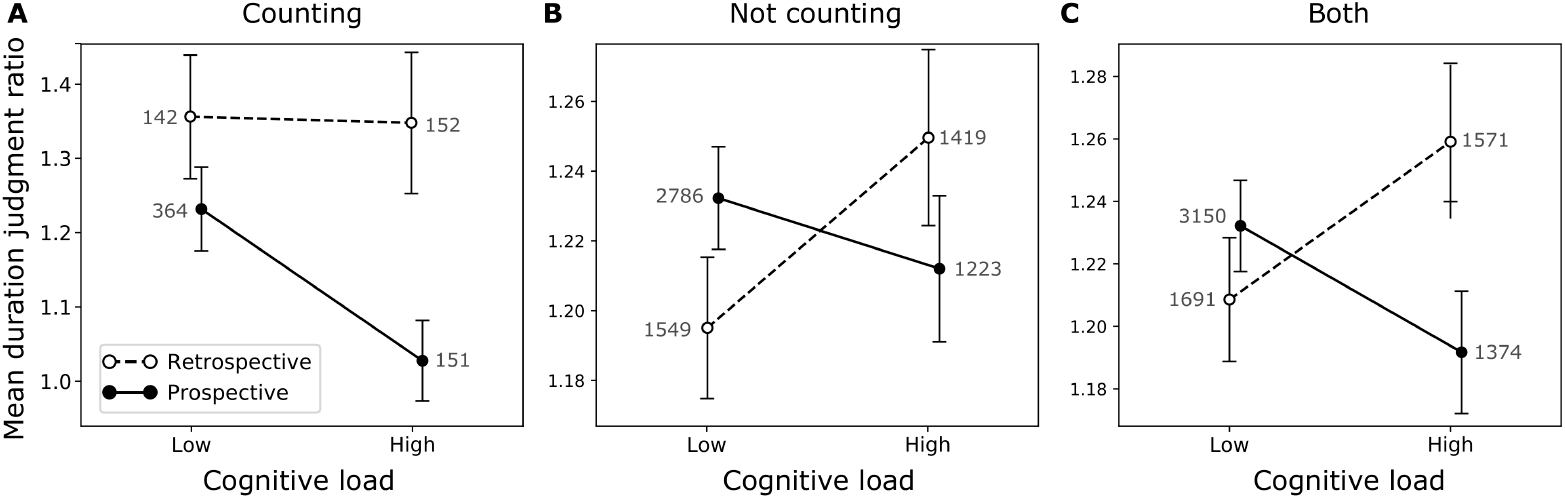
Counting vs not-counting. Interaction in human duration reports tasks (pro/retrospective) and cognitive load for participants that reported they were counting (**A**), not counting (**B**), or both cases together (**C**), having excluded the rejected trials as described in methods.

**S3 Fig.**
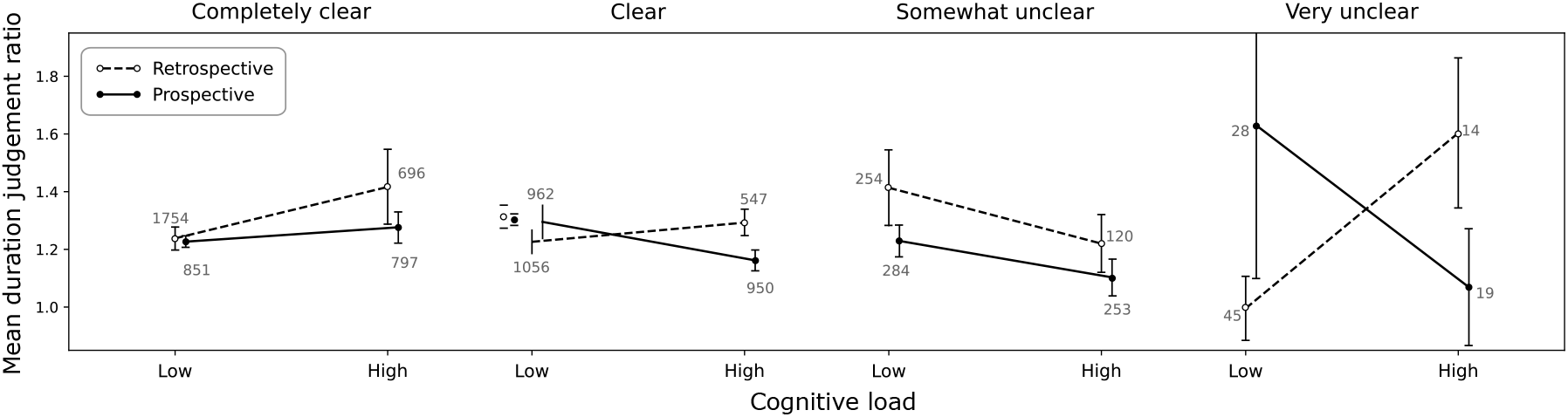
Level of understanding of the task. Interaction in human duration reports tasks (pro/retrospective) and cognitive load for participants depending on how clear they reported they found the initial instructions of task, having excluded the rejected trials as described in methods.

**S4 Fig.**
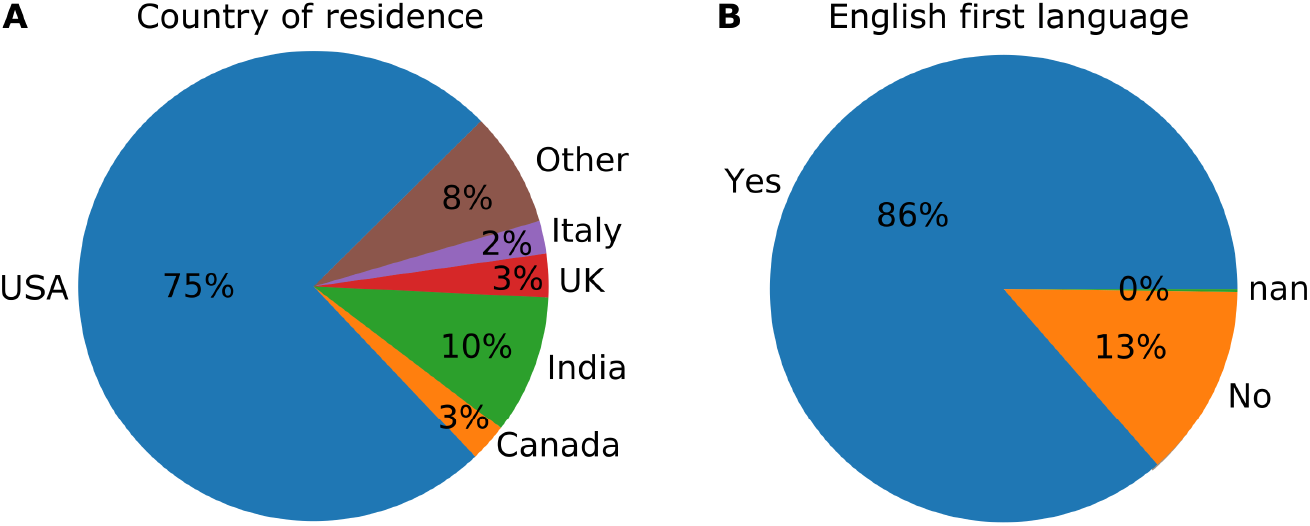
Demographics of the human participants. This includes geographic location and first language.

**S5 Fig.**
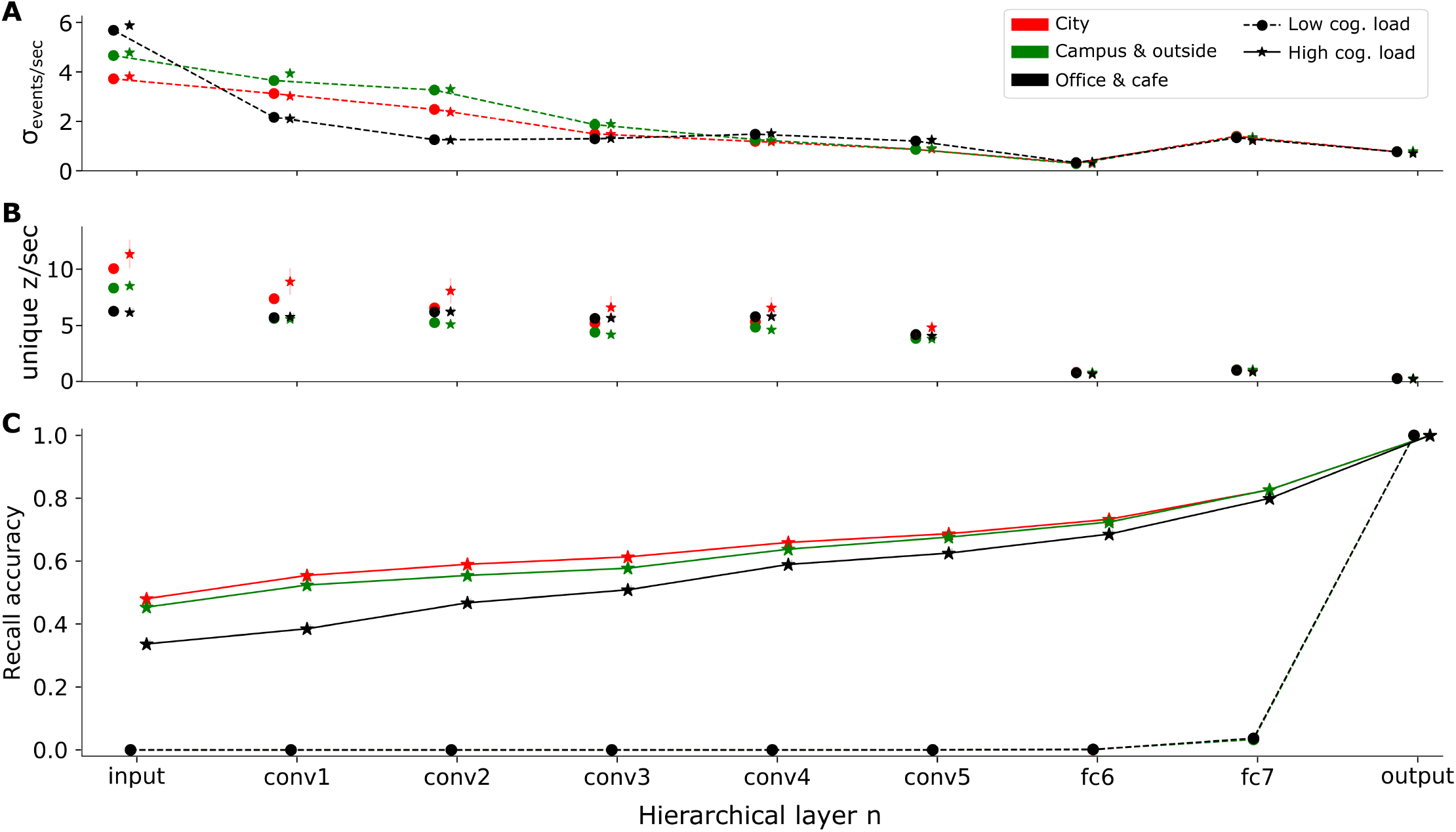
Statistical properties of the model related to episodic memory recall, per scene type and cognitive load level. The episodic memory model shows unique patterns for the 3 different scene types, although no single feature is clearly suitable by itself or superior to alternative solutions to directly select the network layers conv1, conv3 and fc7 as the source of information when estimating time in the corresponding scene types, as shown in Results. **A:** Standard deviation of recalled events per second. Regardless of cognitive load, lower hierarchical layers show far greater variability and, thus, are more affected by the prior of the overall number of events usually recorded. **B:** Number of unique components (different types of events) captured in single episodes. Here, there is a similar overall pattern with A, although more individual components are recorded in cases of high cognitive load. In addition, when comparing different scene types, the unique number of captured components is negatively correlated with the standard deviation of the recalled events. **C:** Accuracy of recalls measured as the number of episodic nodes (events) recalled with components assigned divided by the number of all events recalled in each episode. In cases of low cognitive load only very high-level events can be recalled accurately and estimation is based almost entirely on prior knowledge.

**S6 Fig.**
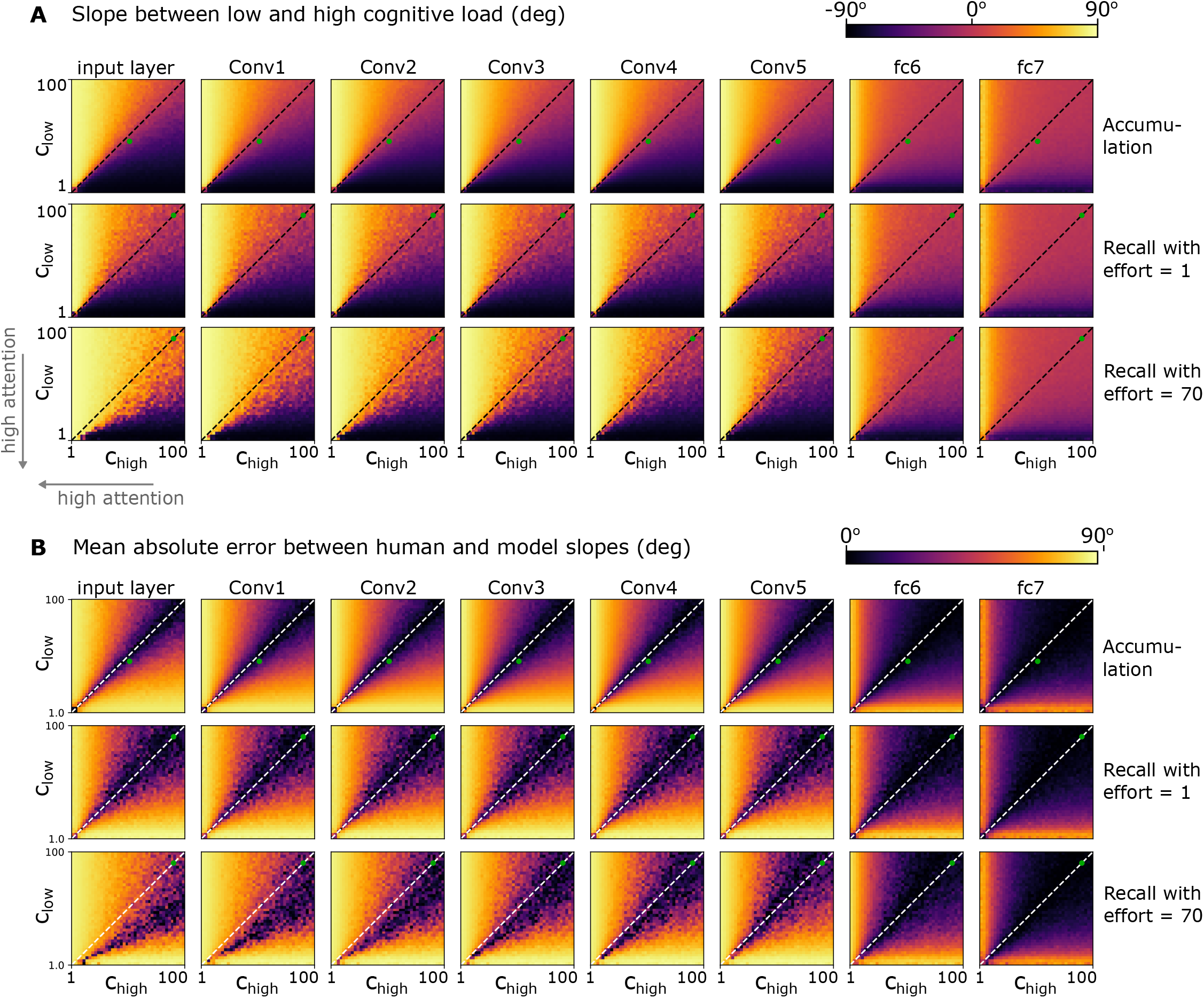
Effects of attention on event accumulation and recall per layer, for the prospective and retrospective tasks respectively. **A**: Slope between the number of accumulated events using low- and high-load hyper parameters. This figure illustrates the slope instead of the absolute number of events in the two task variants, as the absolute number of events is normalized when accumulations are mapped to seconds via linear regression. **B**: The error between the resulting slopes of the computational model and the slopes resulting from human time judgements. The green dots represent the points in the parameter space that were selected and used in Results. This figure shows that in event accumulation and recall with low effort, small differences between *c*_low_ and *c*_high_ are enough to replicate human behaviour. However, under a high amount of recall effort, attention needs to be either significantly lower, in which case human results can be replicated even when *c*_low_ = *c*_high_, or *c*_low_ should become significantly lower.

**S7 Fig.**
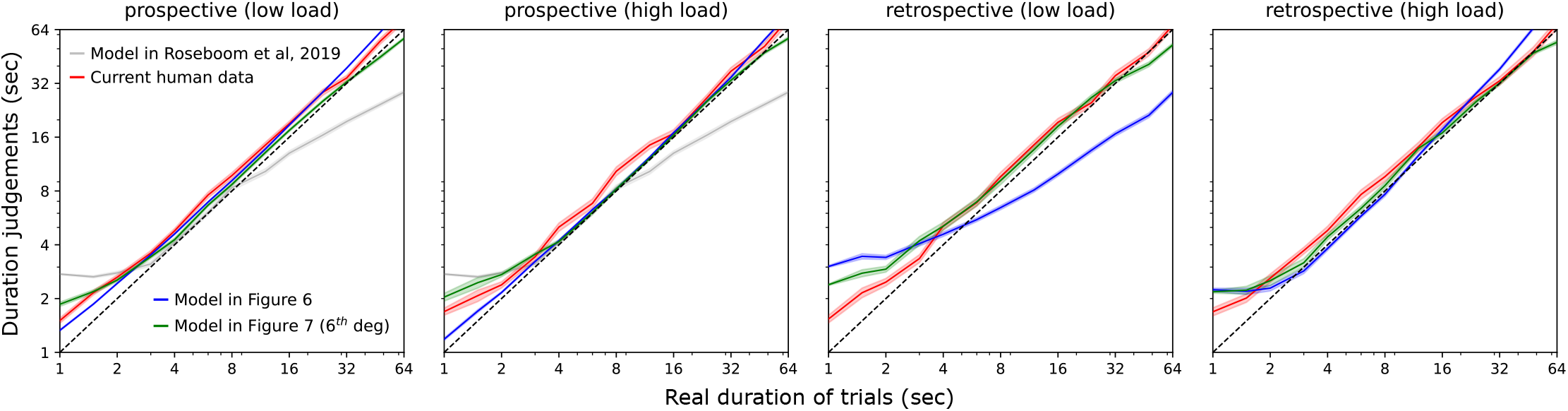
Model comparison. Human duration judgements versus the different versions of the computational model proposed here, as well as the model in [4] that uses “static” frames, for the four variants of the task. The coloured lines and shades areas represent the mean and standard error of the mean respectively.

**S8 Fig.**
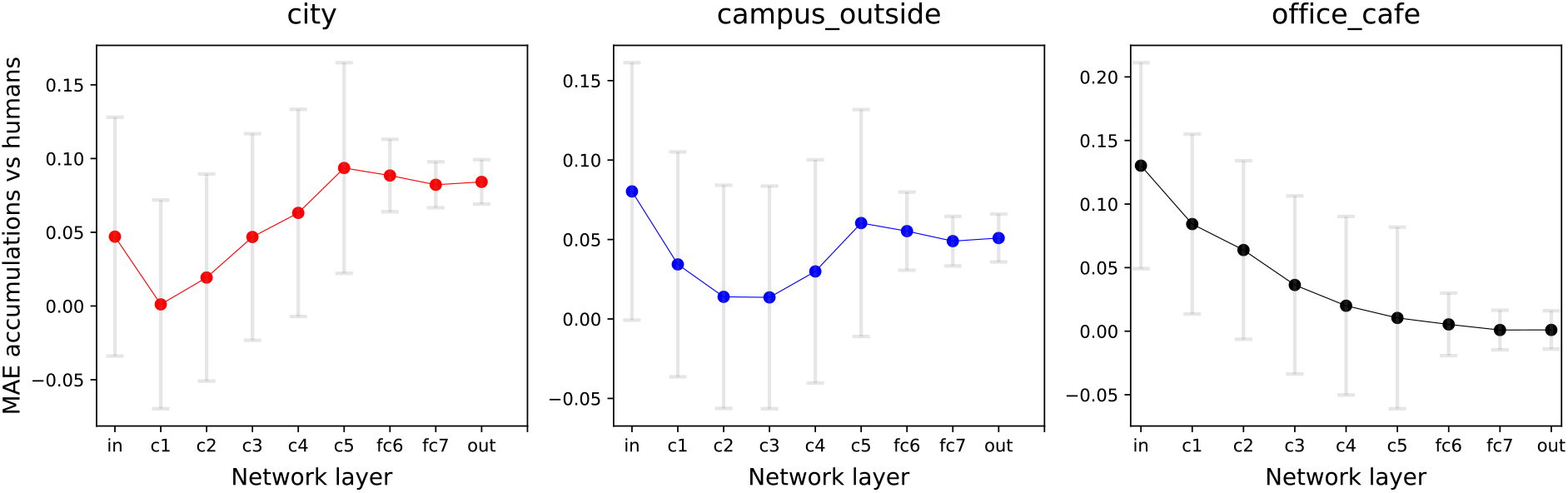
Recalled events per second versus human reports. Mean absolute error between the slope of recalled events per second for low and high cognitive load versus the slope of humans mean duration judgement ratio for low and high cognitive load. Human reports have been segmented per scene type while the graphs compare this error for all hierarchical layers of the neural network. The error bars show standard error of the mean.

**S1 Table.** Probability distributions and corresponding parameters associated with each of the two memory types. Semantic in green and episodic in orange at time *t*.

| Model | Distributions | Associated trainable parameters |
| --- | --- | --- |
| Generative | $P(z_t^n \vert z_t^{n+1}, z_{t-1}^n, e_\tau^n) = \text{Cat}(\mathbf{A}_t)$ | $\mathbf{A}_t(z_t^{n+1}, z_{t-1}^n, e_\tau^n), \mathbf{K}_t^n, \mathbf{V} = \{e_0^0, e_1^0, \dots, e_0^1, \dots\}$ |
| | $P(x_t^n \vert z_t^n) = N(x_t^n \vert \mu_{t,i}^n, \Sigma_{t,i}^n, z_t^n == i)$ | $\mu_{t,i}^n, \Sigma_{t,i}^n$ |
| Inference | $P(x_t^n \vert x_t^{n-1}) = f_{\theta_n}(x_{t-1}^{n-1})$ | $\theta_n$ |
| | $P(z_t^n \vert x_t^n) = \text{argmax}_{i \in \mathbf{K}_t^n} [N(x_t^n \vert \mu_{t,i}^n, \Sigma_{t,i}^n)]$ | $\mu_{t,i}^n, \sigma_{t,i}^n, \mathbf{K}_t^n$ |
| Episodic recall | $P_{\text{evidence}}(e_\tau^n) = \mathbf{W}(e_\tau^n)$ | $\mathbf{V} = \{e_0^0, e_1^0, \dots, e_0^1, \dots\}, \mathbf{W}(e_\tau^n)$ |
| | $P_{\text{chil}}(z_{k,t}^n) = N(\mu_{\text{ch},k}^n, \sigma_{\text{ch},k}^n)$ | $\mu_{\text{ch},k}^n, \sigma_{\text{ch},k}^n$ |
| | $P_{\text{chil}}(n) = N(\mu_{\text{ch}}^n, \sigma_{\text{ch}}^n)$ | $\mu_{\text{ch}}^n, \sigma_{\text{ch}}^n$ |

**S2 Table.** Parameters of the model that can be trained recursively in real-time, along with the training method used.

| Symbol | Parameter description | Training method |
| --- | --- | --- |
| $A_t()$ | Transition probabilities table | Event counter |
| $K_t^n$ | Number of components per layer | Splitting and merging Eq. 6-7 |
| $\mathbf{V}$ | All episodic nodes | Episodic formation Eq. 8 |
| $\mu_{t,i}^n, \Sigma_{t,i}^n$ | Statistics of Gaussian component $i$ | Kalman filter with short-term plasticity Eq. 2-3 |
| $\theta_n$ | Neural network weights and biases | Back-propagation and gradient descent |
| $\mathbf{W}(e_\tau^n)$ | Weights of episodic nodes | Assignment of $P_{\text{evidence}}(x_t^n)$ value when event originally recorded. |
| $\mu_{\text{ch},k}^n, \sigma_{\text{ch},k}^n$ | Number of children nodes for component $k$ | Calculated recursively after every new episodic node. |
| $\mu_{\text{ch}}^n, \sigma_{\text{ch}}^n$ | Number of children nodes for layer $n$ | |

**S3 Table.** Comparison of the proposed single-layer models using the Akaike information criterion (AIC). All listed models were fitted to the real video durations of the same trials that were used for testing. Testing was performed with either real (objective) video durations or with human subjective duration judgements.

| Model | vs human reports | vs real durations | No. of parameters |
| --- | --- | --- | --- |
| Linear (single layer) | 1115 | -1613 | 2 |
| Polynomial 2<sup>nd</sup> deg | 1067 | -4961 | 3 |
| Polynomial 4<sup>th</sup> deg | 635 | -4997 | 5 |
| Polynomial 6<sup>th</sup> deg | **417** | **-5361** | 7 |
| Polynomial 8<sup>th</sup> deg | 1854 | -35 | 9 |

**S4 Table.** Glossary of terms and notation. This glossary includes all notation except the parameters found in S2 Table.

| Notation | Description |
| --- | --- |
| $n \in \{1, 2, \dots\}$ | Index of hierarchical layer. |
| $t$ | Physical time (simulation time step). |
| $\tau$ | Subjective time defined as an index of episodic events. |
| $x_t^0$ | Random variable (continuous) that represents sensory information, in this case frames of a video. This variable follows a mixture of Gaussians. |
| $x_t^n$ | Random variable (continuous) that represents activity of the layer $n$ of the feedforward neural network used, at time $t$. This variable follows a mixture of Gaussians. |
| $\xi_t^n$ | Error between predicted and observed $x_t^n$. |
| $z_t^n$ | Indicative (discrete) variable that represents which Gaussian component is used at time $t$. |
| $e_\tau^n$ | The τ-th event recorded in episodic memory. |
| $\mu_{t,k}^n, \Sigma_{t,k}^n$ | The short-term parameters of the $k$-th Gaussian component. |
| $\tilde{\mu}_{t,k}^n, \tilde{\Sigma}_{t,k}^n$ | The baseline (long-term) parameters of the $k$-th Gaussian component. |
| $K$ | Kalman gain. |
| $Q, R$ | Free noise parameters of Kalman filter. |
| $k_{\text{back}}$ | Model parameter, represents how quickly short-term parameters of a Gaussian component converge back to their baseline. |
| $k_{\text{base}}$ | Model parameter, represents how quickly the baseline parameters adapt to new information. |
| $D_{JS}(\cdot)$ | Jensen-Shannon divergence. |
| $D_{JS}(\cdot)$ | Kullback–Leibler divergence. |
| $T_{\text{merge}}$ | Threshold constant that defines when two Gaussian components merge. |
| $T_{\text{split}}$ | Threshold constant that defines when a Gaussian component splits into two (short-term and baseline parameters become independent). |
| $T_t^n$ | Threshold variable describing the tolerated level of surprise. |
| $\text{Tr}(e)$ | A tree defined as a subset of adjacent recorded episodic events with root node $e$. |
| $P_{\text{chil.}}(z_{k,t}^n)$ | Prior belief of number of children nodes in layer $n - 1$ when the parent node is known (and classified as $z_{k,t}^n$). |
| $P_{\text{chil.}}(n)$ | Prior belief of number of children nodes per recalled node in layer $n$. |
| $P_{\text{evidence}}$ | Equivalent to $P(x_t^n \vert z_{t-1}^n, z_t^{n+1}, e_\tau^n)$. |
| $f(\cdot)$ | Function that maps event accumulations to human readable units of time (e.g. seconds). |
| $\tau_{\text{start}}^n$ | The number of episodic nodes in a layer $n$ at the beginning of an episode that is defined by the tree $\text{Tr}(e^{n^2})$, where $n^2 > n$. |
| $\tau_{\text{end}}^n$ | Same as before, for nodes at the end of an episode. |
| $\tilde{\tau}_{\text{tree}}^n$ | The number of nodes in each layer $n$ of a tree $\text{Tr}(e)$. |
| $D^n$ | Number of simulation iterations since the last time the threshold $T_t^n$ was reset. |
| $c$ | Decaying time constant for threshold $T_t^n$. Used to model the effect of attention. |
| $\epsilon$ | Recall effort. |
| $N(\cdot)$ | Gaussian distribution. |
| $Bern(\cdot)$ | Bernoulli distribution. |

## Acknowledgements

This work was supported by the European Union Future and Emerging Technologies grant (GA:641100) TIMESTORM—Mind and Time: Investigation of the Temporal Traits of Human-Machine Convergence and the Sackler Centre for Consciousness Science. The authors would like to thank Karl Friston for insightful comments on the theoretical model presented here and Chris Bird for comments on a previous version of the manuscript. This research has been approved by the Science and Technology Cross-Schools Research Ethics Committee (C-REC).

## S1 Appendix

### Analysis of the role of effort in memory recall

The final question we investigated is whether, according to our model, the amount of effort that humans commit to recall episodic memories affects their re-experienced sense of duration for these events. Recall effort here is defined as a mechanism that increases the chances of recorded salient events to be retrieved and used as branches of the tree structure shown in Figure 4 (black nodes). The more effort is devoted, the more particular components 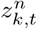 will be part of the recalled experience. Hence, more specific priors 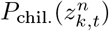 will be used in the recall process over the general prior *P*_chil_. (*n*), to estimate the number of children nodes per recalled node, and deeper in the tree structure 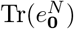 (see Figure 4). To assess how this mechanism affects the rate of recalling events, we ran a series of simulations over the complete set of human trials, where the system received the same video frames as humans and then it produced memory recalls for *ϵ* ∈ {1, 2,…, 70}. As with all simulation results presented here, 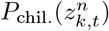 was approximated as the average number of children nodes of the component 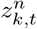 in a single given trial, since only one trial was shown to each subject. In contrast, *P*_chil_. (*n*) was obtained from the distribution of children nodes *n* − 1 for all components in layer *n*, after running simulations for 3300 frames including all scene types.

Figure 9A illustrates the resulting mean values and distributions for all layers of the network. These were 1.2 ± 0.59 children nodes for the convolutional layers (*conv1-5*) and 2.9 ± 2.67 nodes for the fully connected layers (*fc6-7* and *output*), more than twice the values of the former type of layers.

**Fig 9.**
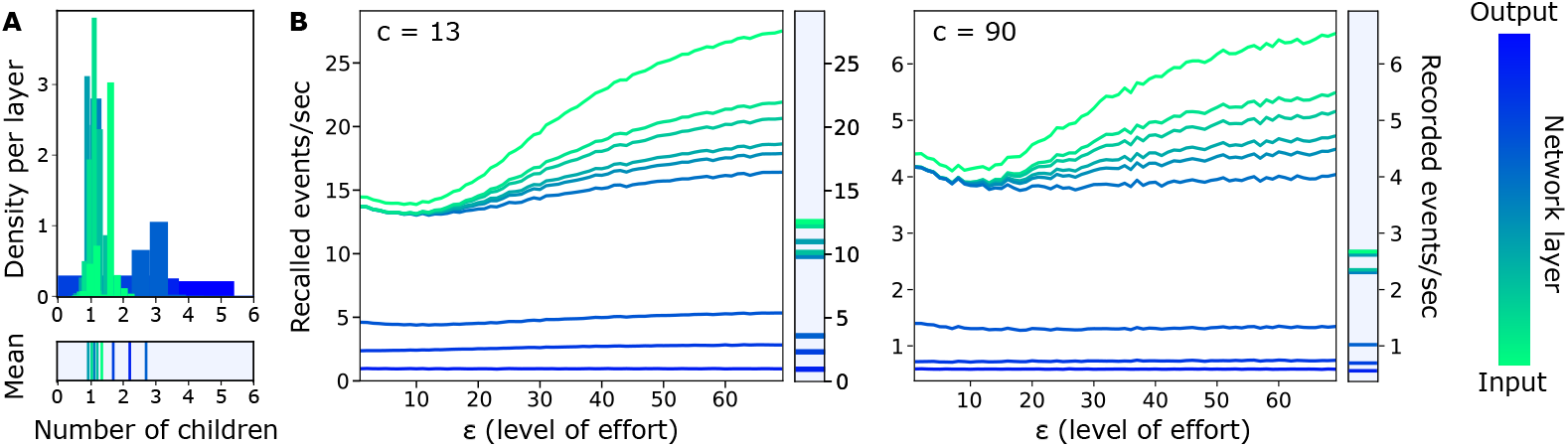
**A:** Distribution of children nodes recorded per layer over all components of this layer, used to calculate *P*_chil_. (*n*). **B:** Rate of recalling events in different layers of the model and for different levels of effort *ϵ*, defined as the number of recall attempts. The coloured lines represent the mean. Standard error was not included in this plot as it overlaps with the mean.

The overall effect of *ϵ* in the rate of recalling events can be seen in Figure 9B. As expected, the layers that are more sensitive to changes in *ϵ* are the lower convolutional ones, since recall is a recursive process that originates from the top (most contextual) layer and has a cumulative effect as it proceeds. For these low layers, high values of *ϵ* resulted in higher number of recalled events, and therefore longer time estimations. Interestingly, this relation was not monotonic and it was maintained for very different levels of attention (e.g. *c* = {13, 90}).

#### Further analysis

Figure 9B illustrates that the relation between effort and accumulated events over time is consistent across different levels of attention and it is non-monotonic. An explanation for this complex behaviour can be summarised as follows. In high layers, where a small number of events occurs in a single episode leaving minimal prior knowledge for individual components 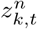, the distribution 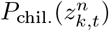 is closer to the real number of recorded events. That is, when trying to recall how many salient events happened during a memory, the richer this memory is in high-level contextual information, the more biased the recall will be by the actual experience. On the other hand, the estimated 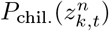 has large variance in low layers, since individual components are more likely to be used multiple times. Indeed, when *ϵ* = 1, only the ~ 5%of the overall recalled events corresponded to nodes from the original episode with particular components 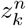 (black nodes in Figure 4), while when *ϵ* = 70, the overall accuracy increased to more than ~ 57%. In addition, the rate of recalled events showed an initial tendency to approach the real number of recorded events, indicated by the bars on the right-hand side of the sub-figures in Figure 9.B. This changed when *ϵ* >7, where the first components of the convolutional layers were recalled, leading to a massive increase in recalled events in these layers.

#### Relation to attention

To further examine how recall effort interacts with attention, and affects the difference between the model’s prospective and retrospective duration judgments, we ran simulations where low and high cognitive load trials were represented by different values of the attention decay time constant *c*, used in Eq (13). Low values of *c* cause the surprise threshold to decay faster, hence denoting higher attention. Additionally, *ϵ* was set to 1 in trials with low cognitive load while, in the opposite case, it was either also 1 or raised to 70. Clearly, when *c*_low_ = *c*_high_ and if there is no effort difference, the rate of recalling events, and therefore duration judgements, is identical in both cases. When attention is considered higher in low cognitive load cases, i.e. *c*_low_ < *c*_high_, the slope between duration judgements resembles the one found in human prospective judgements shown in Figure 5.B. The opposite slope (found in human retrospective judgements) can be achieved when higher effort is used in high cognitive load trials. This interplay between attention and recall effort was robust for a wide range of values of *c*. In fact, the effect of recall effort was significantly more visible in low layers of the network revealing the mechanism that leads to the different patterns of Figure 6.

For more information and a visualization of the model’s behaviour, see S6 Fig.

## S2 Appendix

### Model training process

In the algorithm that follows, pll, phl, rll and rhl correspond to the four variants of the current experiment, i.e. prospective / retrospective and low / high cognitive load respectively, *X_i_* is a matrix that contains accumulated events for *i* ∈ {pll, phl, rll, rhl}, *MDJR_i_* represents the mean duration judgement ratio in human trials for *i, y* a vector that contains the video durations of all trials and MSE stands for mean squared error.

#### Algorithm 2: Complete model training

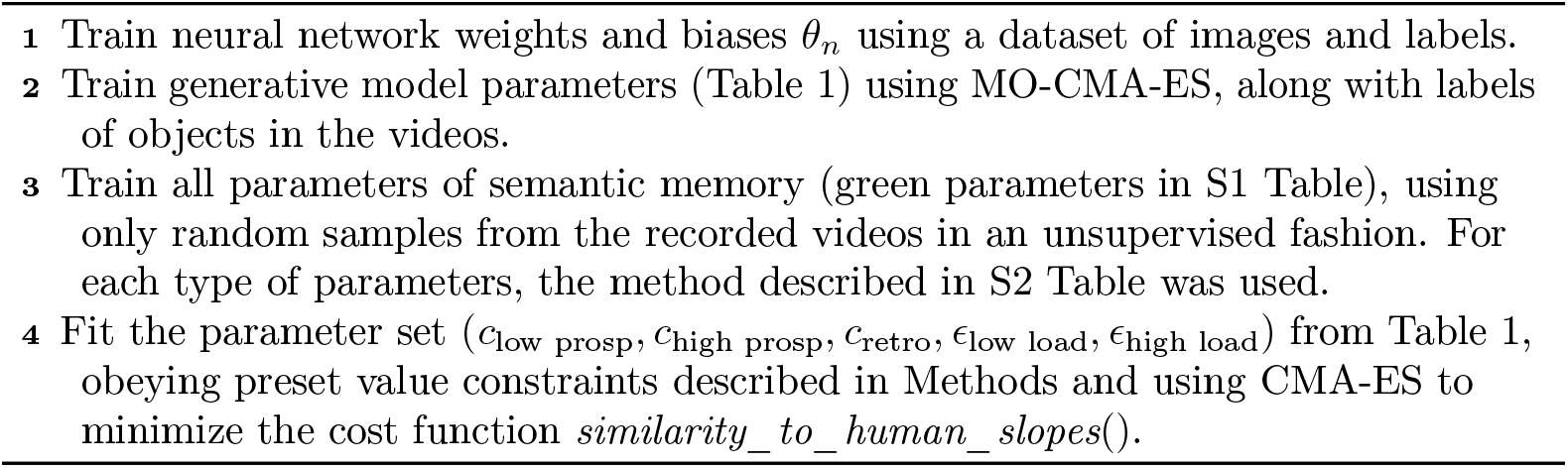

#### Algorithm 3: Cost function for free parameter optimization

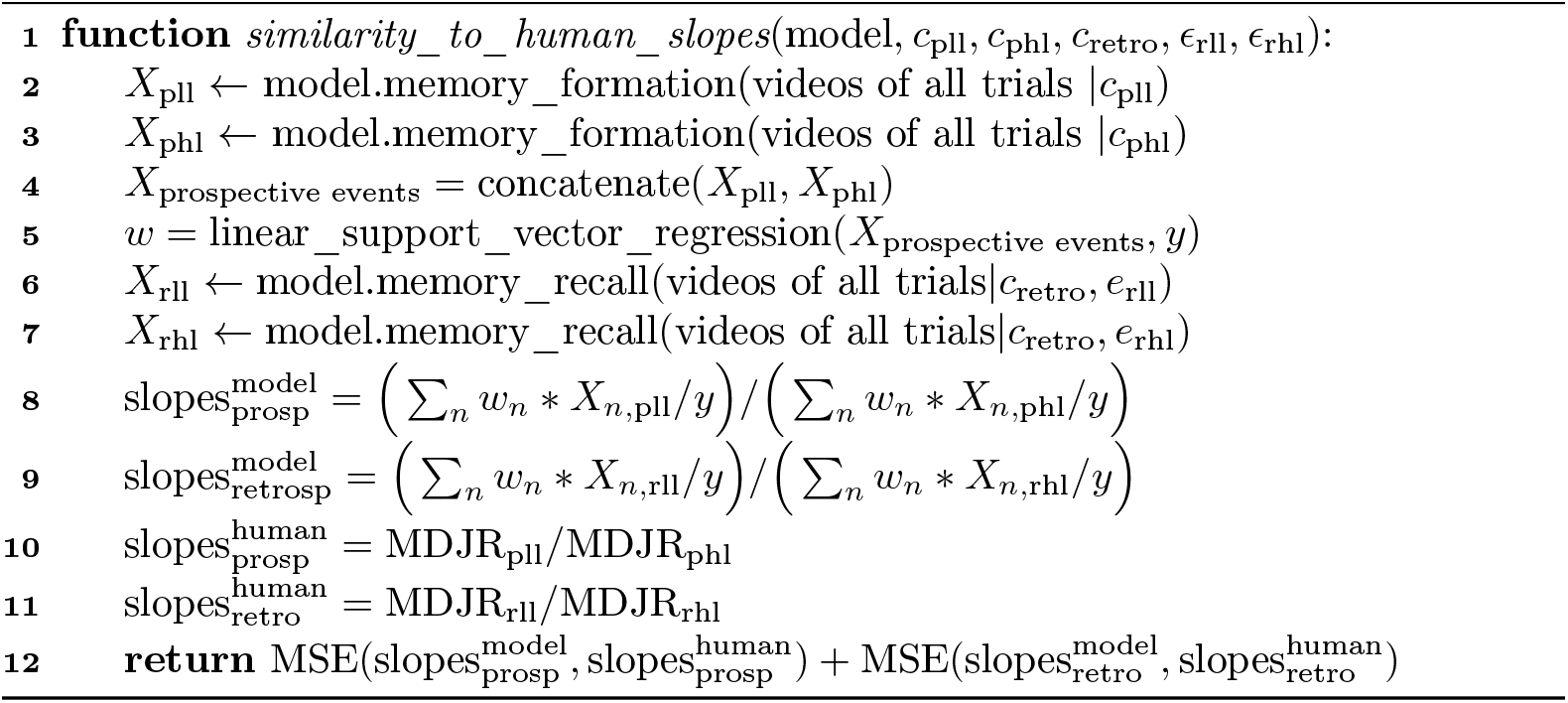

## Footnotes

1 All notation is defined again in the supplementary S4 Table.

2 Human dataset: https://github.com/zfountas/prospective-retrospective-data

3 Source code of the model: https://github.com/zfountas/prospective-retrospective-model

